# Dynamic and decay kinetics of H3 variants in live cells reveal the pivotal role of HIRA/NSD2 in maintaining the distinct H3.3 specific chromatin landscape

**DOI:** 10.1101/2024.05.21.595158

**Authors:** Vishal Nehru, David Ball, Abhishek Mukherjee, Daisuke Kurotaki, Ajay Chitnis, Tatiana S. Karpova, Keiko Ozato

## Abstract

The incorporation of variant histone H3.3 into the genome is tightly linked with transcriptional activity, yet its precise regulatory mechanisms remain elusive. Traditional methods like Chromatin Immunoprecipitation offer static views of H3.3 distribution, lacking dynamic insights. Here, using the SNAP tag system, we employed Fluorescence Recovery After Photobleaching (FRAP) and live-cell imaging to investigate H3.3 mobility and decay kinetics in live mouse embryonic fibroblast cells. Our focus on interferon-induced transcriptional activation revealed rapid H3.3 exchange, indicative of its transcriptional regulatory role. Transcription inhibition hindered H3.3 mobility, emphasizing its involvement in transcription. Additionally, we probed into turnover dynamics(decay) of H3.1-SNAP and H3.3-SNAP variants, uncovering differential decay rates influenced by transcriptional activity and histone modifiers such as NSD2 and HIRA. Live-cell imaging showed faster decay of H3.3 compared to H3.1, further exacerbated upon NSD2/HIRA loss. Notably, HIRA and NSD2, regulators of H3.3 dynamics, proved crucial for both H3.3 mobility and decay, underscoring their pivotal role. These findings deepen our understanding of epigenetic regulation, emphasizing the dynamic nature of histone turnover in cellular function and its implications for disease pathogenesis. Taken together, this study sheds light on the dynamic behavior of H3.3 and its regulatory mechanisms, providing valuable insights into epigenetic regulation in cellular processes and disease contexts.

## Introduction

The histone variant H3.3 is actively expressed throughout the cell cycle and deposited alongside transcription in a manner independent of replication(1). Genome-wide analyses using ChIP-seq technique have unveiled the enrichment of H3.3 within actively transcribed genes, including promoters proximal to transcription start sites (TSSs), as well as within gene bodies and transcription end sites (TESs) across mammalian and Drosophila cells (2–6). The deposition of H3.3 is induced in specific genes upon transcriptional activation(7,8). Additionally, H3.3 is found at telomeres and pericentric heterochromatin, illustrating its diverse localization and functions beyond transcriptional regulation (5,9).

Several methods have been employed to investigate the stability of H3.3 binding within the nucleosome, its dynamic chromatin exchange and turnover(10–14). A conventional method to assess histone stability, exchange and turnover is by exposing cells to radioactive amino acids and tracking the exchange of newly synthesized histones with radiolabeled preexisting histone pool (SILAC)(13).

This method has yielded valuable insights into the dynamic behavior and turnover of histones (13). It has been noted that transcription processes significantly regulate this dynamic exchange and turnover, underscoring the intricate relationship between transcription, histone exchange, and turnover (15). Additionally, during transcription, nucleosomes are displaced and replaced by non-nucleosomal elements, a process that remains complex due to the potential reuse of recycled H3.3 (16). Notably, the deposition of H3.3 is impeded in pericentric heterochromatin by ATRX, while replication influences deposition and gap-filling mechanisms (17). Therefore, investigating global dynamics in live cell conditions, particularly utilizing non-cancerous mammalian cells, is crucial. In this regard, Fluorescence Recovery After Photobleaching (FRAP) emerges as an excellent method.

In this study, using SNAP tag system, we investigated the dynamic behavior of global and newly synthesized H3.3 in live cells and explored the impact of HIRA and NSD2 on this dynamic turnover(15). Furthermore, by combining with photobleaching technique FRAP (Fluorescence Recovery After Photobleaching), allowed us to analyze diffusion coefficients, exchange rates of SNAP tagged proteins, and determine the proportions of mobile and immobile fractions in vivo(16,17). H3.1-SNAP (WT), H3.3-SNAP (HIRA cKO), and H3.3-SNAP (NSD2-KO) displayed restricted entry into the bleached area even after extended periods of bleaching, suggesting their immobilization within nucleosomes. Conversely, H3.3-SNAP (WT) promptly infiltrated the bleached region, facilitating the restoration of fluorescence. Furthermore, our results indicate that cell cycle progression proceeded without discernible alterations in kinetics. Histone turnover, particularly decay, stands as a cornerstone process in chromatin dynamics, intricately shaping gene expression patterns and cellular functions. Delving into the decay dynamics of H3.1 and H3.3 histone variants has unveiled crucial insights into the orchestration of chromatin structure and function (16,41). Our investigation not only underscores the distinct roles of H3.1 and H3.3 in chromatin regulation but also delineates their differential decay kinetics. While H3.1 tends to associate with constitutively active chromatin regions, H3.3 preferentially deposits at actively transcribed sites, showcasing nuanced decay patterns that underscore their unique regulatory mechanisms and functional significance in chromatin dynamics.

Moreover, our study delves into the intricate interplay between transcriptional activity and histone turnover. Transcriptional inhibition prompts a delayed decay of H3.3, unveiling the dynamic responsiveness of histone turnover to changes in transcriptional programs and emphasizing the dynamic nature of chromatin remodeling(18). Further elucidating the regulatory landscape, our investigation highlights the pivotal roles of histone-modifying enzymes, such as NSD2, and chaperone protein, HIRA, in governing histone turnover dynamics. Loss of NSD2 notably impacts the decay of H3.3, suggesting a role for NSD2-mediated histone methylation in H3.3 stability, while loss of HIRA delays the decay of both H3.1 and H3.3, implicating a broader role for HIRA in histone turnover regulation beyond its canonical function in H3.3 deposition. Understanding the dynamics of histone turnover holds profound implications for epigenetic regulation and disease pathogenesis. Dysregulation of histone turnover has been implicated in various pathological conditions, including cancer and neurodegenerative disorders. In summary, our study offers invaluable insights into the dynamic behavior of H3.3 in live mouse fibroblast cells, shedding light on its continuous interaction with chromatin and reaffirming its direct involvement in transcriptional processes. The delineation of distinct binding kinetics and the exploration of its temporal dynamics under transcriptional arrest enrich our understanding of H3.3’s regulatory mechanisms governing gene expression control.

## Materials and Methods

### Cells

Wild-type (WT), and HIRA^fl/fl^ Cre-ER^T^ MEF cells were prepared from E12.5 embryos and maintained in Dulbecco’s minimal essential medium (DMEM) supplemented with 10% fetal bovine serum (R&D systems) and 1% antibiotics (Gibco). NSD2^-/-^ MEF cells was a kind gift from Dr. Kiyoe Ura (Chiba University, Japan). To conditionally knock-out HIRA, cells were treated with Tamoxifen(4uM) for 4 days and knock-out was confirmed by qPCR and western blot. Cells were stimulated with mouse recombinant IFNβ (100units/ml, PBL Interferon Source) for indicated time-points. Flavopiridol (100nm, Sigma-Aldrich) and Actinomycin D (1µg/ml, Sigma-Aldrich) or vehicle (DMSO) was added to the cells with and without IFNβ depending on the experimental design.

### DNA constructs and Transduction

The cDNAs for H3.1-SNAP-3XHA and H3.3-SNAP-3XHA were cloned sequentially into pMSCV-puro retroviral vectors obtained from Addgene (Plasmid 68469). Briefly, the constructed vectors were first transfected into the RetroPack PT67 Cell Line (Takara-631510) and transduction to introduce the constructs into the target cells using the procedures previously described (19).

### Antibodies

The primary antibodies were purchased from the following retailers: HA (ab9110), H3K36me2(ab9049), H2B (ab64165) were from Abcam, and H3K36me3(9763) from Cell Signaling Technology. Horseradish peroxidase-conjugated secondary antibodies (ECL anti-rabbit IgG, HRP-linked species-specific whole antibody (from Donkey) and ECL anti-mouse IgG, HRP-linked species-specific whole antibody (from sheep) were from Amersham-GE healthcare.

### Immunocytochemistry

For immunofluorescence (IF), coverslips were first coated with poly-L-Lysine (Millipore-Sigma) for 1h at room temperature, rinsed and UV sterilized. Cells were seeded and fixed with 4% paraformaldehyde in PBS, pH 7.4 for 10-15 minutes at room temperature, washed three times with ice-cold PBS. The cells were then permeabilized for 10 min in PBS containing 0.25% Triton X-100, washed, and blocked with 10% normal goat serum in 0.1% PBS-Tween (PBST) for 1h, washed and incubated with primary antibodies diluted in 1%BSA in PBST for 1h at room temperature. The primary antibodies were washed three times with PBST for 5 min and cells were then incubated with secondary antibodies for 1h in the dark. As above, antibodies were washed three times in PBST, and cells were counterstained with either DAPI (1:2000) or Hoechst (1:250) for two minutes, washed again and mounted using ProLong Diamond Antifade Mountant, (P36961).

### Reverse transcription and qRT-PCR

For total RNA isolation, the ZYMO research kit (R1054) was utilized, followed by cDNA synthesis using Superscript II (Invitrogen). The cDNA samples were subjected to qRT-PCR using the SYBR Green Master Mix (Applied Biosystems) and the ABI Prism 7500 sequence detection system (Applied Biosystems). *Hprt* and *Gapdh* were used as reference genes to normalize the transcript levels.

### Protein Extraction, Analysis by SDS-PAGE, and Western Blotting

Histone preparation was carried out following the established protocol (22). Nuclear extracts were resolved on NuPAGE Tris 4-12% polyacrylamide gels in MES buffer (Invitrogen) and subsequently transferred to nitrocellulose membranes (Protran). For immunodetection, primary antibodies including rabbit anti-HA (ab9110, Abcam) and anti-H2B (64165, mAbcam) were utilized. Horseradish peroxidase-conjugated secondary antibodies (Amersham-GE healthcare) were employed for the detection of primary antibody-bound proteins. Chemiluminescent detection was performed using SuperSignal West Pico Plus chemiluminescent detection kits (Thermo Fisher), and the resulting chemiluminescent signal was captured using the Azure Biosystems c600 imaging system.

### RNA-seq

RNA-seq analysis was conducted following the standard Illumina protocol. Total RNA was extracted from Mouse Embryonic Fibroblasts (WT-MEFs) and NSD2 knockout MEFs using the ZYMO RNA kit (R1054). Library construction was performed using the Illumina stranded mRNA-seq kit, following the manufacturer’s instructions. A total of 500 ng of RNA was used for library preparation. The pooled libraries were subsequently sequenced using an Illumina NextSeq 500 sequencer. To align the RNA-seq Fastq files to the UCSC mm10 genome, we employed Bowtie2 aligner(20). Read counts per gene were generated using HTSeq-count, Salmon suite and differential expression analysis was conducted using edgeR(21). For visualization on the IGV genome browser, normalized bigWig files were generated using deepTools(22). RNA-seq were performed two times.

### Real time live cell imaging and Microscopy

For the preexisting fraction, MEF cells expressing H3.1-SNAP and H3.3-SNAP were initially plated in Nunc Lab-Tek II two-well chambered coverglass (ThermoFisher Scientific). Prior to live cell imaging, the H3.1-SNAP and H3.3-SNAP were labeled with SNAP-Cell 505-Star (New England Biolabs, S9103S) for a duration of 30 minutes. This was followed by two washes with PBS and one wash with DMEM. The cells were then incubated for an additional 30 minutes with DMEM (containing FBS but no antibiotics), and a final wash with DMEM was performed.

After the above-mentioned steps, live cell imaging was conducted using a spinning disk microscope (Nikon) with an incubator chamber stage. Timelapse image acquisition was done using a 20X (NA=0.75) objective and an Andor DU-897 X-7308 camera on a Nikon Spinning Disk Confocal microscope. 16-bit images, without binning, were acquired in the blue and green channels with lasers having excitation wavelengths of 405 nm and 488 nm respectively, with an exposure time of 200ms. The acquisition window size was 512 x 512 pixels, but that corresponded to 240.39 µm x 240.39 µm for H3.1-SNAP-3XHA (WT) and H3.3-SNAP-3XHA (WT) samples; and to 359.80 x 359.80 µm for H3.1-SNAP-3XHA (NSD2^-/-^) and H3.3-SNAP-3XHA (NSD2^-/-^) samples.

For the new pool/fraction, MEF cells expressing H3.1-SNAP and H3.3-SNAP were first blocked using SNAP-Cell Block (New England Biolabs, S9106S) to inhibit the preexisting fractions/populations of H3.1-SNAP-3XHA and H3.3-SNAP-3XHA. This was followed by a chase period of approximately ∼4 hours, and then the cells were washed twice with PBS and once with DMEM (with FBS). Subsequently, live cell imaging was performed using a spinning disk microscope (Nikon) for a duration of ∼50 hours using a 20x objectives and a filter set with an excitation maximum at 504 nm and an emission maximum at 532 nm to capture 3D images without z-sectioning. Images were acquired every 10 minutes for the first hour, and then every 30-60 minutes for up to 500 minutes. Following image acquisition, the images were analyzed using Fiji Image J (23) and Imaris software (Oxford Instruments).

### Quantification of H3.1 and H3.3 fluorescence intensities

Fluorescence intensities from two-dimensional timelapse movies, consisting of DAPI (blue) and H3.1-SNAP or H3.3-SNAP (green) channels, were quantified using FIJI (ImageJ)(23). Both channels of each timelapse video were split and converted into 8-bits. Backgrounds of both channels were subtracted using the “Subtract Background” process in FIJI to remove background noise and autofluorescence. “Bleach Correction” was then applied to the DAPI channel using the “Histogram Matching” method to correct for any possible photobleaching due to imaging over long durations (∼ 48-50 hours). The “Otsu” thresholding method was then used to create binary nuclei masks from the bleach corrected DAPI channels for all timepoints in each experimental condition. The “Fill Holes” technique was then used on the masks to maximize coverage of nuclei area. The resultant masks were then “AND” processed with the green channel (H3.1 or H3.3) to ensure that only fluorescence intensity from within the nuclei were extracted and processed further. Mean signal intensities of the green channels for each timepoint were then extracted. The resultant signal intensities at each timepoint were then normalized with that at the start of image acquisition (t = 0 h) and plotted in MATLAB (2021b) and t^1/2^ is the time taken by a signal to decrease by half of its original value.

### Fluorescence recovery after photobleaching (FRAP)

FRAP analysis was conducted following the established protocol (34). Imaging was performed using a Zeiss confocal laser-scanning microscope 780 equipped with a 45 mW Ar laser. Prior to FRAP, live SNAP cells were labeled with SNAP-505-Star dye and incubated for 30 minutes. Subsequently, the cells were washed several times with PBS and DMEM and maintained at a temperature of 37°C and 5% CO2 on a heated stage incubator (ASI 400, Nevtek, Burnsville, VA).

To perform the bleaching, a single image was captured with 0.3% laser power, a pinhole aperture of 4, fast scan mode, and 7x zoom. A selected area with a radius of 0.5μm, approximately 25 pixels (indicated in yellow color), was then bleached by scanning twice with 100% laser power for approximately 0.2 seconds. After a 15-30 second recovery period, subsequent images were acquired using the same imaging settings as the pre-bleach image.

The intensity of the bleached area relative to the entire nucleus was measured using Fiji ImageJ2 software, after subtracting the background signal. The intensity values were then compared to the pre-bleached image to determine the relative changes in intensity. Additionally, SNAP cells labeled with 505-Star labelled dye were fixed with 4% paraformaldehyde for 30 minutes to establish baseline autofluorescence levels for further analysis.

During the live-cell imaging period, nuclei exhibited movements, primarily rotations about the z-axis. Nuclei that rotated about the xy-axis were excluded from the analysis, while the bleached area remained easily identifiable. The association and dissociation kinetics were analyzed based on the assumption that tagged histones diffuse within the nuclei over minutes. FRAP experiments were performed on a minimum of 15-20 independent cells, and the data were averaged to generate a single FRAP curve. The FRAP data were then analyzed and fitted to a mathematical binding model (34). These experiments were performed three times.

## Results

### Cell Lines Expressing H3.1-SNAP-3XHA and H3.3-SNAP-3XHA

To gain insights into the dynamics of H3.3 and examine the influence of NSD2 and HIRA on H3.3 dynamics, we used mouse embryonic fibroblast (MEF) cells as a model system. These cells were transduced with a retroviral preparation containing an expression plasmid (pMSCV-puro) encoding histones H3.1-SNAP-3XHA and H3.3-SNAP-3XHA cDNAs. The expression plasmid was designed to be driven by the PGK promoter, ensuring a constitutive and near-endogenous expression of these histone variants in the transduced cells (Fig. S1A-C).

The incorporation of the SNAP tag allowed efficient labeling and visualization of the total or preexisting/global as well as the newly synthesized populations or fractions of H3.1-SNAP and H3.3-SNAP histones in live cells, enabling the analysis of their dynamics, decay and behavior during transcription and within the nucleus (Fig. S1B-C)(24,25).

Immunostaining of H3.1-SNAP-3XHA and H3.3-SNAP-3XHA for the preexisting fraction revealed distinct staining patterns. H3.1-SNAP exhibited localization to both euchromatin and heterochromatin, while H3.3-SNAP showed a stronger presence in euchromatin regions (Fig. S1B). Furthermore, the expression of the SNAP-tagged histone variants was confirmed by immunodetection using an anti-HA antibody, confirming successful expression in the transduced MEF cells (Fig. S1C).

To further examine the functional interplay between HIRA and NSD2 in the dynamics of H3.1 and H3.3, H3.1-SNAP-3XHA and H3.3-SNAP-3XHA were transduced into HIRA conditional knockout (HIRA cKO) and NSD2 straight knockout (NSD2 KO) MEF cells. RNA sequencing (RNA-seq) analysis was performed to assess the transcriptome changes demonstrated by the downregulation of NSD2 in the NSD2-KO cells compared to wild-type (WT) cells (Fig. S1D). In the case of HIRA cKO cells, cells were first treated with Tamoxifen (5μM) for 4 days to induce HIRA knockout, followed by qRT-PCR analysis. The expression of HIRA mRNA was significantly reduced when normalized to the *GAPDH* gene. The downregulation was further confirmed by immunodetection using an anti-HIRA (WC-119 clone) antibody, with H2B serving as a loading control (Fig. S1E, F and G).

To ensure that the introduction of the SNAP-tagged histone variants did not interfere with cell cycle progression or their incorporation into chromatin, we examined the cell cycle progression of the transduced cells expressing H3.1-SNAP-3XHA and H3.3-SNAP-3XHA. Interestingly, our analysis revealed that the transduced cells exhibited similar kinetics during cell cycle progression as the untransduced cells (data not shown).

These observations suggest that the presence of the SNAP tag on the histone variants did not disrupt the transcription and cell cycle. It also indicates that the incorporation of H3.1-SNAP-3XHA and H3.3-SNAP-3XHA into chromatin and their subsequent behavior during cell division were not significantly affected by the presence of SNAP tag, supporting the suitability in our experimental system for studying the dynamics and decay kinetics of H3.1 and H3.3 variants.

### Fluorescence recovery after photobleaching (FRAP) elucidates the molecular dynamics of H3 variants and insight into distinct fractions

To delve deeper into the molecular dynamics more specifically mobility within the nucleus of these H3 variants, we utilized the Fluorescence Recovery After Photobleaching (FRAP) technique on live mouse fibroblast cells. Specifically, we utilized live MEF cells stably expressing H3.1-SNAP, H3.3-SNAP, H3.3-SNAP (Hira cKO), and H3.3-SNAP (NSD2 KO), which were labeled with a green, fluorescent SNAP-Cell permeable 505-Star dye to visualize preexisting fractions in addition to newly synthesized pool of H3 variants. By selectively photobleaching a small region of interest (ROI) within the nucleus at t_0_, we continuously monitored the intensity within this region every 10 minutes for a duration of up to 500 minutes. In theory, if all molecules exhibit complete mobility (i.e., freely diffusible, or capable of dissociating and rebind within 60 minutes), the intensity in the bleached area, relative to that in the entire nucleus, would rapidly reach its maximum within the initial 60 minutes and remain constant thereafter. Conversely, if all molecules are immobile (i.e., bound for more than 60 minutes), the recovery after 60 minutes would be either slow or rapid, depending largely on the rates of dissociation and reassociation. If a fraction of molecules is mobile, this fraction would exhibit recovery by t_120min_, and any subsequent increase would be contingent upon dissociation and association rates as calculated in (Figs. S5A and S5B).

Fig. 1A illustrates images from a representative experiment, while the results from multiple experiments are depicted in Figs. B-E, and fraction quantification is presented in Fig. F. Importantly, these experimental procedures were conducted without adverse effects on cell viability, as evidenced by the occurrence of mitosis and subsequent cell division during the live-imaging period.

**Figure 1.**
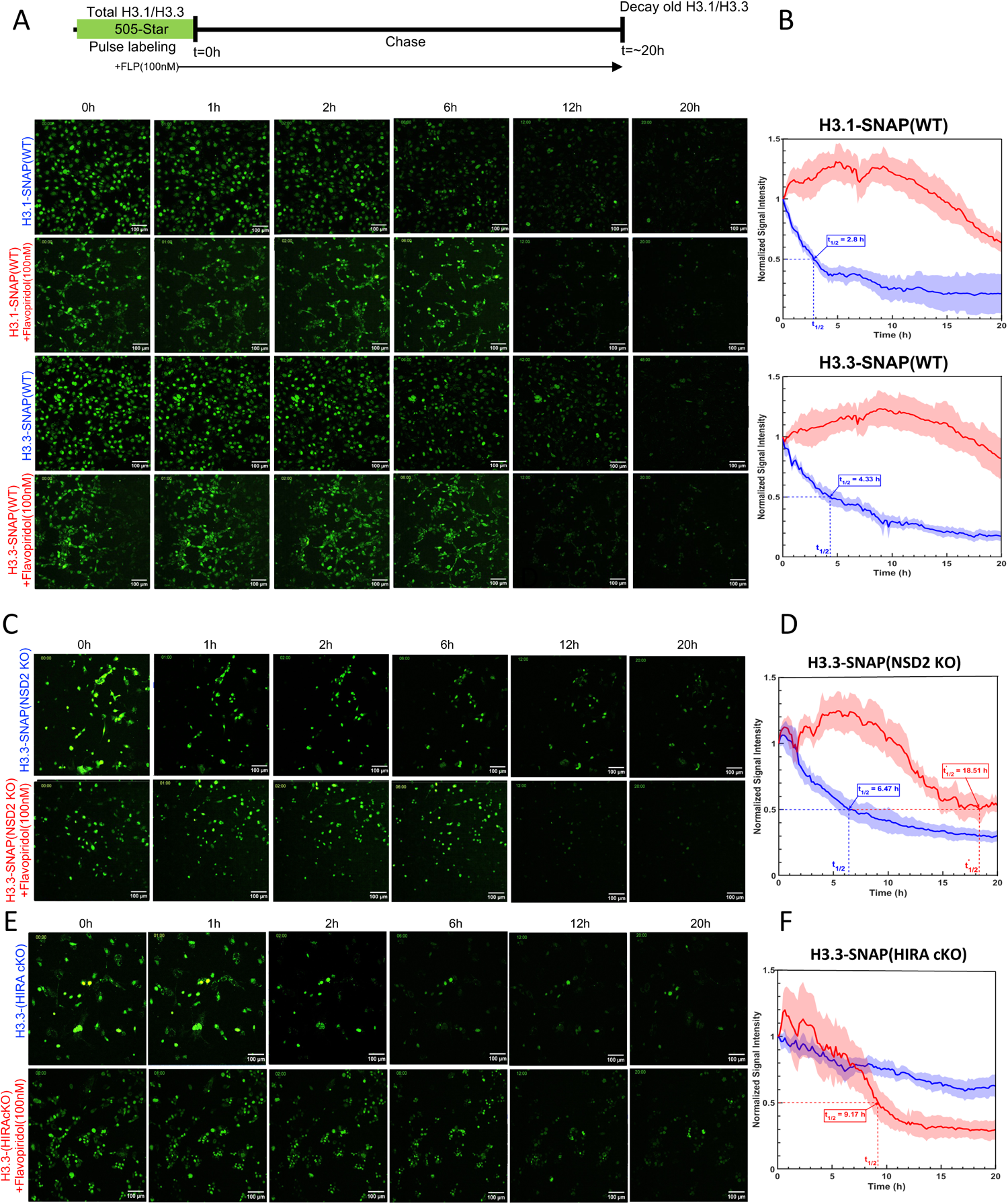
Measuring the kinetics of H3 variants elucidates the molecular dynamics of H3 variants and insight into distinct fractions. **A**. FRAP approach: Cells expressing H3.1-SNAP-3XHA(WT) and H3.3-SNAP-3XHA(WT) were labeled with SNAP-Cell compatible 505-Star dye, and a small region of the nucleus was bleached. Confocal imaging was performed every 10 minutes for 1 hour, followed by imaging every 30-60 minutes for up to 500 minutes. The same procedure was conducted for H3.3-SNAP-3XHA (HIRA cKO) and H3.3-SNAP(NSD2KO) cells, Scale bars, 10 μm. **B-E**. FRAP curves: Compared to H3.1-SNAP, H3.3-SNAP is mobile within the nucleus and loss of HIRA, NSD2 affects H3.3 mobility. Interpretation: If H3.1-SNAP and/or H3.3-SNAP were mobile, they repopulated the bleached region, resulting in a decrease in overall fluorescence in the nuclei. Conversely, if they were immobile, the bleached region remained unchanged. Interpretation. The intensity of fluorescence is reduced by bleaching to a “baseline” level, which was determined using fixed cells. Subsequently, the fluorescence recovers as unbleached molecules enter the bleached region. The rate and extent of recovery are dependent on the contribution of different kinetic fractions. Here, a “mobile” fraction diffuses into the bleached region before the first image is collected after bleaching. Then, a “rapid” fraction becomes initially dominant, followed by a “slow” fraction that progressively becomes more important. There is also a “very slow” fraction, but its entry is too slow to be monitored. **F**. The percentage of the population and the association half-life (t_1/2_) of these different fractions were determined using data from panels B-E and summarized in supplementary table ST1A.

Data collected from H3.3-SNAP expressing cells revealed distinct recovery kinetics, fractions compared to H3.1-SNAP. H3.3-SNAP exhibited a rapid rate of fluorescence recovery, as indicated by relatively mobile-fast fraction of (50%), mobile-slow fraction of (28%) and immobile fraction of (23%), while H3.1-SNAP displayed slower recovery kinetics (immobile-fraction of 74%, mobile-fast of 21% and mobile-slow of (5%) indicative of distinct recovery kinetics (Fig. 1B-E and quantitation provided in F; Table ST1A; Supplementary Figures. S5A and S5B). This suggests compared to H3.1-SNAP, most of H3.3-SNAP molecules were mobile (dynamically exchanging with chromatin) at any given time and relatively less bound(immobile) fraction quantitation provided in Fig.1F.

Interestingly, the dynamics of H3.3-SNAP were further altered in the absence of Hira and/or NSD2. H3.3-SNAP (Hira cKO) exhibited significantly reduced mobility compared to wild-type H3.3-SNAP(WT) of immobile or bound fraction (86%), fast exchange or Mobile-Fast (7%) and Mobile-Slow (7%), highlighting the direct role of Hira in regulating H3.3 mobility with the chromatin (Fig. 1E; Table ST1A; Supplementary Figures. S5A and S5B). Similarly, H3.3-SNAP (NSD2 KO) also displayed partial impaired mobility with immobile-fraction (50%), Mobile-Fast (22%) and Mobile-Slow (28%), implicating NSD2 in modulating H3.3 dynamics (Fig. 1D; Table ST1A Supplementary Figures. S5A and S5B). These observable differences in molecular dynamics provide insights into their functional significance. The relatively higher mobility of H3.3 suggests its involvement in dynamic chromatin regions, while the reduced mobility of H3.1 indicates a potential role in more stable chromatin structures. The altered dynamics of H3.3 in the absence of Hira or NSD2 underscore the importance of these factors in regulating H3.3. Together, these findings highlight the intricate interplay between histone variants, chromatin dynamics, and associated regulatory factors.

Furthermore, we also observed distinct characteristics of the newly synthesized fraction of H3.3 compared to the preexisting pool/fractions. The newly synthesized H3.3 fraction exhibited reduced dynamics and a higher propensity to associate with chromatin, indicating greater immobility (60%), Mobile-Fast (29%) and Mobile-Slow (11%) compared to the preexisting or global H3.3 immobile-fraction (23%), Mobile-Fast (50%) and Mobile-Slow (28%) Fig. S2(A-C; Table T2B Supplementary Figures. S5Aand S5B). This further emphasizes the differences in dynamics and chromatin binding between the new and preexisting H3.3 fractions. In addition, activation by interferon-stimulation(IFN-β) altered the newly synthesized H3.3 fraction mobility with immobile-fraction (41%), Mobile-Fast (35%) and Mobile-Slow (24%) compared to that of preexisting or global H3.3 immobile-fraction of (23%), Mobile-Fast of (50%) and Mobile-Slow of (28%) further emphasizing that new pool dynamics is different to that of preexisting fraction Fig. S2(D) (for comparison refer, Supplementary Tables, ST1A and ST1B).

In conclusion, our findings provide significant insights into the molecular dynamics of H3 variants, shedding light on their unique roles in chromatin regulation. Moreover, the presence of a newly synthesized H3.3 fraction with distinct slow mobility highlights the intricate interplay between H3 variants and their functional properties within the chromatin environment.

### Activation of transcription by IFNβ stimulation arrests H3.3 dynamics

Activation of transcriptional machinery is a crucial process for regulating gene expression and controlling various cellular activities (26). One aspect of transcriptional regulation involves the dynamics of histone variants, such as H3.3, which contribute to the modulation of chromatin structure and gene accessibility(27).

To investigate the impact of IFNβ stimulation on H3.3 dynamics, we again employed a live-cell imaging approach coupled with fluorescence recovery after photobleaching (FRAP) analysis. We utilized H3.3-SNAP as a molecular probe to monitor the dynamics of H3.3 in response to IFNβ stimulation. By conducting and monitoring live-cell imaging over a 50-hour period and labeling cells with a SNAP-compatible dye, we were able to observe the fluorescence recovery and evaluate the dynamics of H3.3.

Our results revealed that the activation of transcriptional machinery by IFNβ stimulation(100U/ml) led to a reduction in the mobility of H3.3. Specifically, we observed a significant decrease in the mobility and exchange rate of H3.3 within the chromatin upon IFNβ treatment (Mobile-fast fraction 22%, Mobile-Slow fraction 78%) and no immobile or bound fraction. This suggests that the transcriptional activation induced by IFNβ directly influences the stability and dynamics of H3.3, potentially affecting chromatin structure and gene accessibility (Figs. 2, A, C and E, and Table ST1A; Supplementary Figures S5A and S5B). The new pool fraction dynamics was different from that of global pool (Mobile-fast fraction of 35%, Mobile-Slow fraction of 24% and immobile or bound fraction of 41%) suggesting its dynamics behavior within the chromatin is different from that of the preexisting pool of H3.3 (Fig. S2, Table T2B) To further explore the molecular mechanisms underlying the mobility of H3.3 dynamics upon IFNβ stimulation, we investigated the involvement of H3.3 chaperone, HIRA(Fig.2D). Previous studies have highlighted the role of HIRA in the deposition of H3.3(28). In cells with HIRA conditional knockout (Hira cKO), we observed loss of HIRA exacerbated the arrest of H3.3 mobility even after IFNβ stimulation (Fig.2, D and E). This suggests a potential interplay between HIRA-mediated H3.3 deposition and reinforces the role HIRA in transcriptional regulation of H3.3.

**Figure 2.**
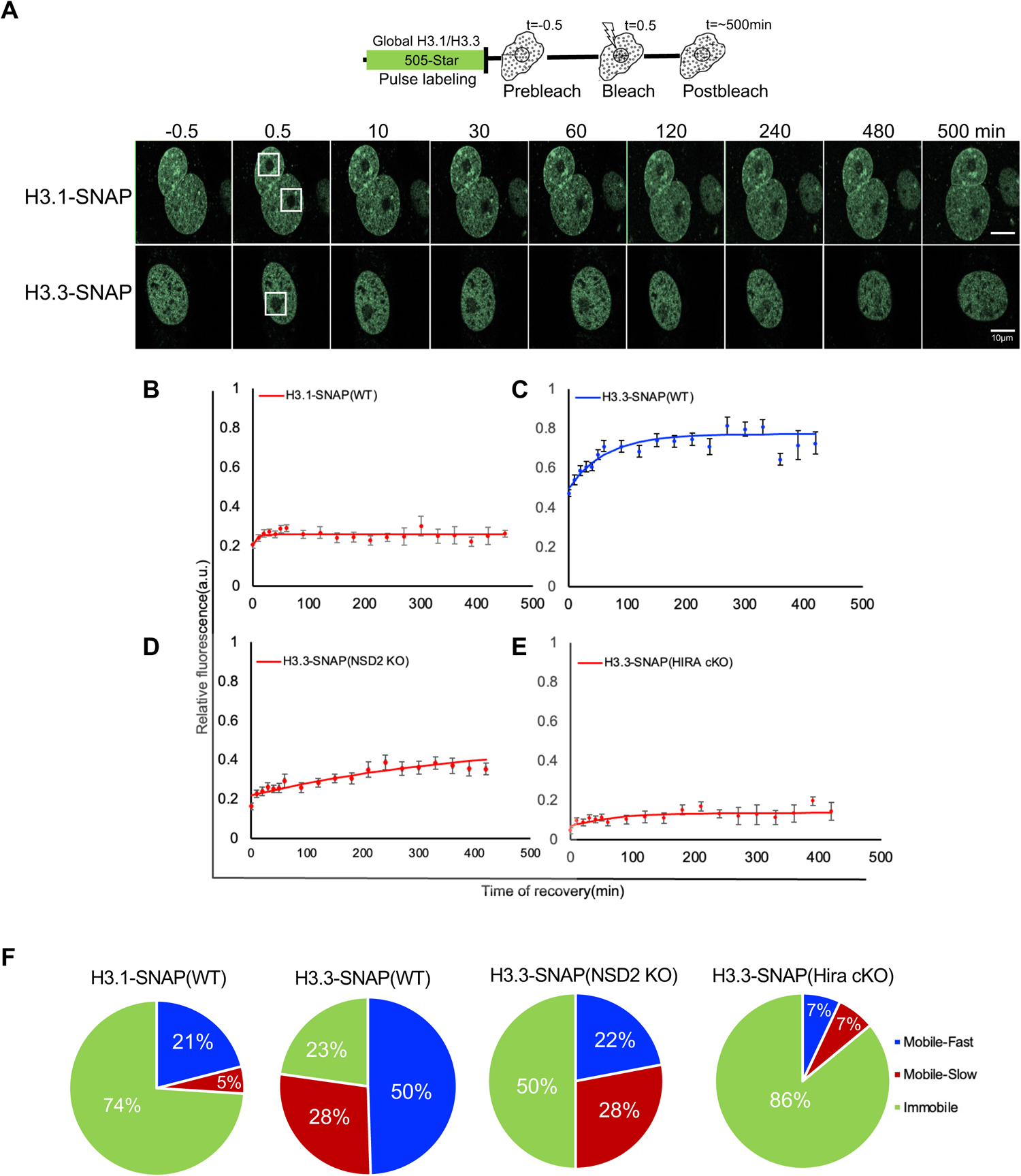
Activation of transcriptional machinery by IFNβ stimulation immobilizes H3.3. **A**. Time-lapse confocal images of a bleached area within a nucleus expressing H3.3-SNAP. Images were taken every 10 minutes for 1 hour and every 30 minutes thereafter. Representative images are shown. Scale bars represent 10 μm. **B-D**. Fluorescence Recovery After Photobleaching (FRAP) curve: The fluorescence intensity in the nuclei decreased due to bleaching, and subsequent recovery depended on the mobility of H3.3-SNAP (IFNβ-), H3.3-SNAP (IFNβ+) and/or H3.3-SNAP-Hira cKO (IFNβ+). The bleached region remained unchanged if the molecules were immobile. The interpretation of the FRAP curve is described, where different kinetic fractions contribute to the recovery process. **E**. Population percentage and associated half-life (t^1/2^) of different kinetic fractions were determined using data from panels B-D and in Supplementary Table ST1A.

Our findings shed light on the intricate relationship between the activation of transcriptional machinery, and H3.3 dynamics. The arrest of H3.3 dynamics upon IFNβ stimulation highlights the significance of transcriptional regulation in modulating chromatin structure and gene expression. Additionally, our results highlight the dynamic nature of H3.3 and its sensitivity to external stimuli. The alteration of H3.3 dynamics by IFNβ suggests that H3.3 plays a crucial role in mediating the cellular response to interferon signaling, potentially influencing the expression of interferon-responsive genes, and modulating immune responses(19).

In conclusion, our study provides compelling evidence that the activation of transcriptional machinery by IFNβ stimulation influences the arrest of H3.3 dynamics. These findings also contribute to our understanding of the intricate molecular mechanisms governing chromatin dynamics and gene regulation. The observed arrest of H3.3 dynamics upon IFNβ stimulation suggests a functional role for H3.3 in the transcriptional regulation associated with interferon signaling and anti-viral immunity(29). The immobilization of H3.3 due to loss of HIRA chaperone may facilitate the stable binding and/or exchange of H3.3 to specific genomic regions, contributing to the establishment of either transcriptionally active or repressive chromatin states(30,31).

### Transcription inhibition alters H3.3 dynamics

To investigate the impact of transcription inhibition on H3.3 dynamics, we tested two transcription inhibitors namely, Actinomycin D and Flavopiridol(32,33). Actinomycin D is a potent inhibitor of RNA polymerase activity, and Flavopiridol is a cyclin-dependent kinase inhibitor known to suppress transcription by inhibiting the activity of CDK9/P-TEFb, a key regulator of RNA polymerase II during elongation stage (33). By treating cells with Actinomycin D and/or Flavopiridol, we aimed to examine how transcription inhibition affects the mobility of H3.3(18,34,35).

We labeled cells expressing H3.3-SNAP with SNAP-compatible 505-star dye (green) and Hoechst stain (blue), enabling live-cell imaging. Initially, we imaged the cells under normal conditions to establish a baseline for H3.3 dynamics. Subsequently, we treated the cells with Actinomycin D and/or Flavopiridol and performed FRAP on live cells over a period of ∼8 hours (∼500 minutes) (Figs. 3 A-D, E: Fig S4.A-C, D-E and Table ST1A and ST2B; Supplementary Figures. S5A and S5B).

**Figure 3.**
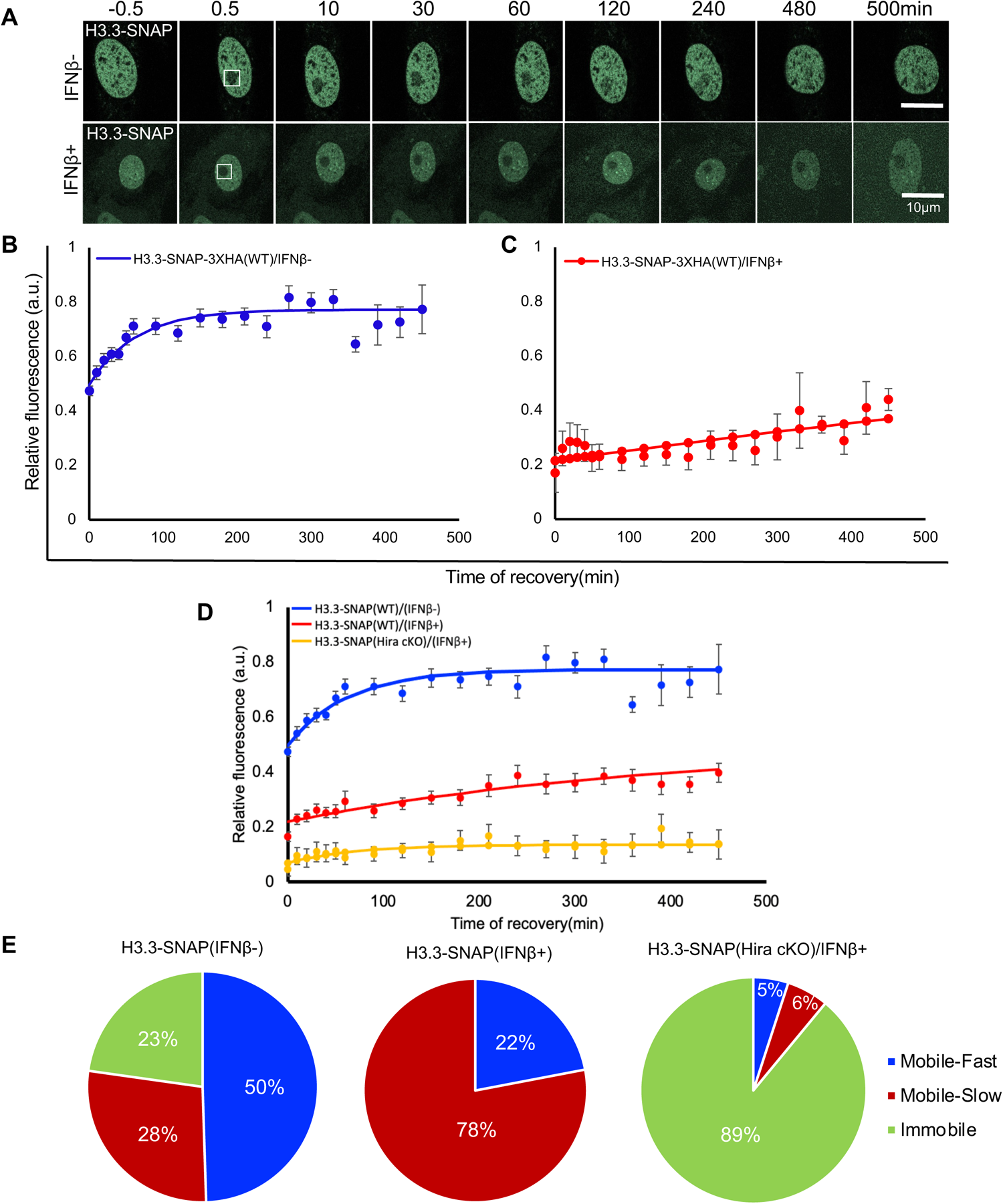
Transcription inhibition impacts the H3.3 dynamics. A. Experimental setup strategy for SNAP Cell 505-star dye labeling of SNAP-tag expressing H3.3-SNAP-3XHA(WT) MEF cells, with and without Actinomycin D(1ug/ml) and Flavopiridol(100nM) and IFNβ. The schematic illustrates the cell/nuclei permeable dye and labeling procedure. B-D. FRAP curves: Cell treated with Actinomycin D and Flavopiridol arrest H3.3 mobility even after IFNβ stimulation reinforcing H3.3 role in transcription couple deposition. E. Population percentage and associated half-life (t^1/2^) of different kinetic fractions were determined using data from panels B-D, Figure. S5A, and Supplementary Table ST1A. Actinomycin D(1ug/ml) and Flavopiridol(100nM) impede H3.3 mobility despite IFNβ treatment.

As observed, in the untreated condition, H3.3 recovery post bleach occurred normally with nearly half (50%) being Mobile Fast fraction, and Mobile Slow (28%) and Immobile Fractions (23%). In contrast, cells treated with Actinomycin D (1µg/ml) showed a marked decrease in H3.3 recovery post bleach, indicating a reduced rate of replacement or removal of H3.3 from chromatin, with Mobile Fast fraction (12%), Mobile Slow fraction (20%), and Immobile fraction (68%).

In addition, cells treated with Flavopiridol (100 nM) displayed a significant reduction in H3.3 recovery compared to the untreated condition. Similar to the effects observed with Actinomycin D, the impaired transcription caused by Flavopiridol treatment resulted in altered H3.3 dynamics. Specifically, we observed a decrease in the Mobile Fast fraction (18%), an increase in the Mobile Slow fraction (21%), and an increase in the Immobile fraction (61%). These findings indicate that the inhibition of p-TEFb hinders the deposition and exchange of H3.3. It also suggests that the process of transcriptional elongation and H3.3 deposition, although linked by NSD2, are executed through partially distinct molecular mechanisms, where HIRA plays a distinct role(18). We next examined the recovery of newly synthesized H3.3 pool and obtained the fractions in the presence of Actinomycin D, of Mobile Fast fraction (33%), the Mobile Slow fraction of (9%), and the Immobile fraction of (58%). Similarly, with Flavopiridol treatment, we observed a Mobile Fast fraction of (20%), a Mobile Slow fraction of (9%), and an Immobile fraction of (71%). These findings suggest that treatment with either Actinomycin D or Flavopiridol affects the dynamics of newly synthesized H3.3, indicating a potential impact on transcriptional processes. These findings highlight the influence of P-TEFb inhibition on H3.3 mobility and its interplay with transcriptional regulation. The distinct effects observed with Actinomycin D and Flavopiridol treatments suggest the involvement of unique molecular mechanisms in transcriptional elongation and H3.3 deposition(36).

These results also demonstrate that inhibiting transcription has a direct impact on the mobility dynamics of H3.3, further supporting the link between transcriptional activity and H3.3. The sensitivity of H3.3 dynamics to changes in transcriptional activity highlights the dynamic nature of H3.3 and its close association with transcriptional processes.

In conclusion, our findings provide insights into the intricate relationship between transcriptional activity and the dynamics of histone H3.3. It also suggests that the molecular mechanisms governing H3.3 dynamics are influenced by transcriptional processes, and any perturbations in transcription can have a profound impact on the dynamics and functional roles of H3.3. The combined effect of transcriptional inhibition and interferon treatment further exacerbates the alteration in H3.3 dynamics highlighting the intricate interplay between transcriptional activity and the dynamics of histone variants, shedding light on the functional implications of H3.3 in gene expression and regulation.

### Altered decay kinetics of H3.3 compared to H3.1 exacerbated by the loss of NSD2/HIRA

With FRAP analysis, we examined the dynamics (mobility) of both H3.1 and H3.3 histone H3 variants. Previous findings have highlighted histone turnover as a crucial regulator of cell type-specific transcription and plasticity in the mammalian brain. This study sheds light on a novel mechanistic role for HIRA (histone cell cycle regulator) and proteasomal degradation-associated histone dynamics in modulating activity-dependent transcription, synaptic connectivity, and behavior uncovering a striking developmental profile of nucleosome occupancy across the lifespan of rodents and humans, with the histone variant H3.3 reaching near-saturating levels throughout the neuronal genome by mid-adolescence. Despite this accumulation, H3.3-containing nucleosomes remain highly dynamic, independent of modifications, thereby governing neuronal- and glial-specific gene expression patterns throughout life (13).

Expanding on this knowledge, we delved into the decay kinetics of H3.1-SNAP and H3.3-SNAP to unravel their longevity within the nucleus. Unlike studies that relied on fixed-cell imaging at various time intervals, we opted for real time tracking of H3.1-SNAP and H3.3-SNAP decay with live-cell imaging spanning for 50 hours. Cells were pulse-labeled with SNAP-compatible 505-star dye (green) and Hoechst stain (blue) at t_0_ and tracked(chased) over the entire duration, revealing intriguing differences in decay rates between H3.1 and H3.3(28). Remarkably, H3.3 exhibited a faster decay compared to H3.1, as evidenced by t_1/2_, indicating the time taken for a signal to decrease by half of its original value, thus suggesting a higher rate of removal of H3.3 from chromatin (Fig. 4A), with quantification provided in (B).

**Figure 4.**
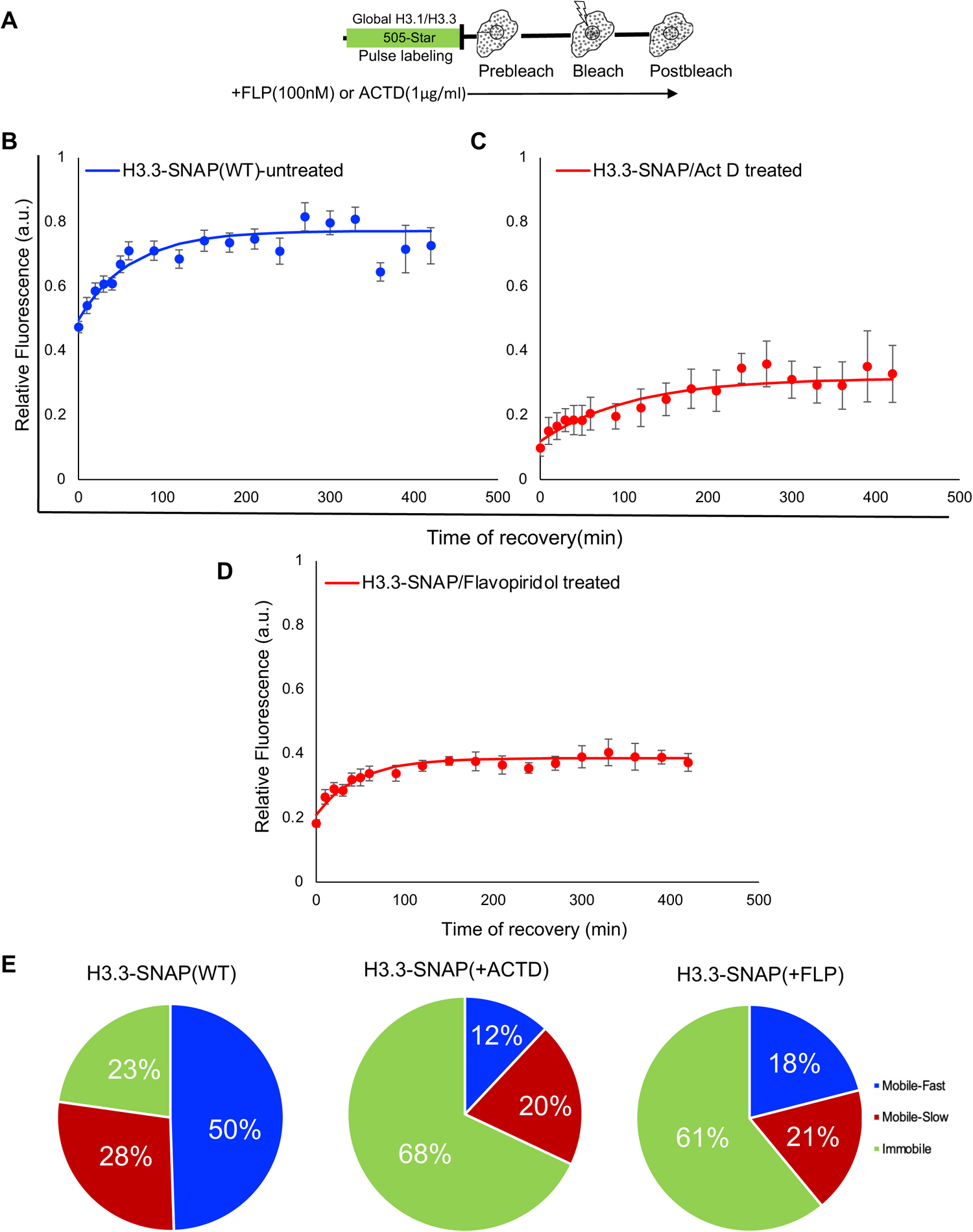
Altered decay kinetics of H3.3 compared to H3.1 exacerbated by the loss of NSD2/HIRA. A. Experimental schematic. Displays the pulse-chase fluorescence intensity of H3.1-SNAP-3XHA (WT) and H3.3-SNAP-3XHA (WT) at different timepoints (0 h, 1 h, 2h, 6h, 12h and 48 h). B. Shows the fluorescence decay curves of H3.1-SNAP-3XHA (WT) (represented by the blue curve) and H3.3-SNAP-3XHA(WT) (represented by the red curve) with t^1/2^. C. Experimental schematic. Displays the pulse-chase fluorescence intensity of H3.3-SNAP-3XHA (WT) and H3.3-SNAP-3XHA (NSD2 KO) at different timepoints (0 h, 1 h, 2h, 6h, 12h and 48 h). D. Shows the fluorescence decay curves of H3.3-SNAP-3XHA (WT) and H3.3-SNAP-3XHA (NSD2 KO) with t^1/2^. E. Experimental schematic. Displays the pulse-chase fluorescence intensity of H3.3-SNAP-3XHA (WT) and H3.3-SNAP-3XHA (HIRA cKO) at different timepoints (0 h, 1 h, 2h, 6h, 12h and 48 h). F. Shows the fluorescence decay curves of H3.3-SNAP-3XHA (WT) and H3.3-SNAP-3XHA (HIRA cKO) with t^1/2^. All scale bars represent 100 µm.

Moreover, previous studies have also highlighted the crucial role of the HIRA chaperone in facilitating H3.3 deposition during transcription(13). We hypothesized and investigated the loss of HIRA and its impact on the decay rates of H3.3 (Fig. 4E), with quantification elucidated in (F)(38). Compared to control, loss of HIRA impacted H3.3 decay kinectics. Furthermore, the role of NSD2 in decay of H3.3 have not been well established despite its mechanistic role being already documented in H3.3 deposition on the ISGs(18).

To address this, we repeated our labeling and performed live-cell imaging using mouse embryonic fibroblast (MEF) cells lacking NSD2 (NSD2KO). Interestingly, our findings provide compelling evidence that the absence of NSD2 further exacerbated the decay(turnover) of H3.3, indicating that NSD2, along with HIRA, plays a critical role in modulating the dynamics and stability of H3.3 within the chromatin landscape (Fig. 4C), with quantification provided in (D). In addition, we also investigated the decay kinetics of H3.1-SNAP (Fig. S8A), with quantification provided in (B). Interestingly, the loss of NSD2/HIRA exhibited an effect on H3.1 decay, with a more pronounced impact observed in HIRA cKO condition. However, the precise mechanism through which the loss of HIRA influences H3.1-SNAP decay remains unclear. It is noteworthy that the HIRA chaperone is known for its specific role in depositing H3.3 onto active chromatin and does not participate in the deposition of H3.1. Thus, understanding the observed impact on H3.1 decay in the absence of NSD2/HIRA presents an intriguing avenue for further investigation.

We next examined the impact of transcriptional inhibition (i.e., Flavopiridol) on the decay of H3.1 and H3.3. To achieve this, cells were initially pulse-labeled with SNAP-compatible 505-star dye (green) and Hoechst stain (blue) at t_0_, followed by treatment with +FLP (100nM), and then tracked over a continuous period of 20 hours. This experimental setup unveiled notable differences in decay rates between H3.1 and H3.3. Surprisingly, H3.3-SNAP demonstrated a delayed decay kinetics as did H3.1-SNAP, indicating an accumulation of H3.1/H3.3 in chromatin (as illustrated in Fig. 5A), with detailed quantification provided in (B). This observation suggests a nuanced relationship between transcriptional activity and the dynamics of histone turnover, warranting further exploration into the underlying mechanisms governing these processes. The observed delayed decay of H3.3 compared to H3.1 upon transcriptional inhibition can be attributed to the distinct roles and regulatory mechanisms governing these histone variants. H3.3 is known to be preferentially incorporated into actively transcribed regions of the genome, where its turnover rate is tightly linked to ongoing transcriptional activity.

**Figure 5.**
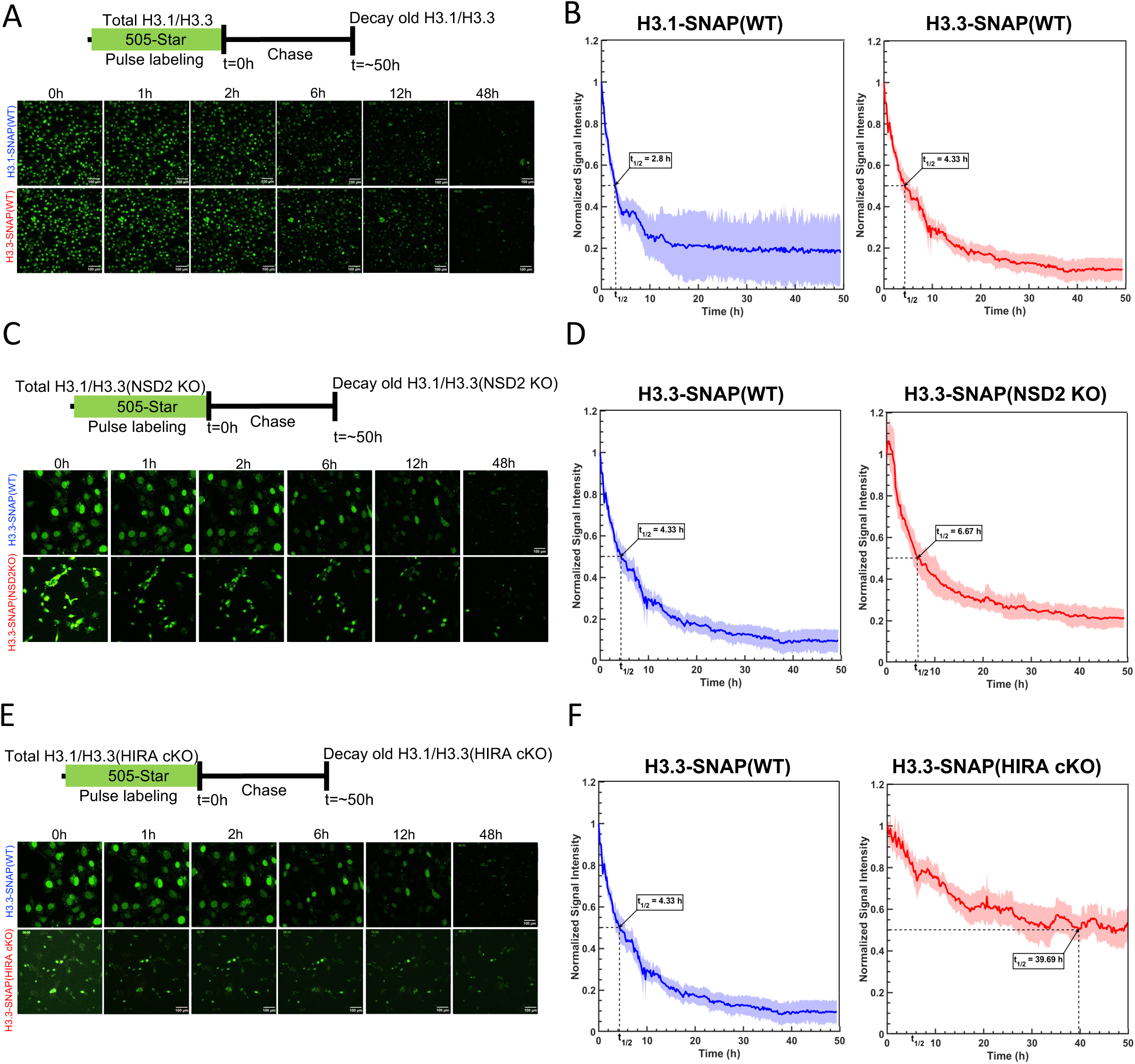
Loss of NSD2/HIRA Combined with Transcription Inhibition Alters H3.3 Decay Kinetics. A-B. Experimental strategy for SNAP Cell 505-star dye labeling of SNAP-tag expressing H3.1-SNAP-3XHA(WT) and H3.3-SNAP-3XHA(WT) MEF cells, with and without Flavopiridol (100 nM). Treatment with Flavopiridol arrests H3.3 decay, as observed by the half-time (t^1/2^) decay kinetics in H3.1-SNAP-3XHA(WT) and H3.3-SNAP-3XHA(WT) MEF cells, determined using data from panels in A and Supplementary Figure S6(A-F). C-F. Similar to B, H3.3-SNAP-3XHA(NSD2KO) and H3.3-SNAP-3XHA (HIRA cKO) MEF cells were treated with and without Flavopiridol and H3.3 decay was measured in terms of half-life (t^1/2^) decay, determined using data from panels C and Supplementary Figures S7(A-F) and S8(A-D). All scale bars represent 100 µm.

Transcriptional inhibition likely disrupts the normal turnover dynamics of H3.3 by halting the deposition of new H3.3 molecules into chromatin at active transcription sites. Consequently, existing H3.3 molecules remain associated with chromatin for an extended period before being replaced, leading to the observed delay in decay kinetics.

In contrast, H3.1, being predominantly associated with constitutively active regions of the genome, may exhibit a more rapid turnover rate even under conditions of transcriptional inhibition. This could be due to the continuous exchange of H3.1 molecules facilitated by general chromatin remodeling processes, independent of ongoing transcriptional activity.

Overall, the differential response of H3.1 and H3.3 to transcriptional inhibition underscores the complex interplay between histone variant-specific deposition mechanisms, transcriptional activity, and chromatin dynamics, highlighting the need for further mechanistic studies to fully elucidate these processes.

Expanding our investigation, we delved into the roles of NSD2 and HIRA in regulating the decay dynamics of both H3.1 and H3.3 histone variants in presence of transcriptional inhibitor. Intriguingly, we found that the loss of NSD2 led to a delay specifically in the decay of H3.3. (as illustrated in Fig. 5C), with detailed quantification provided in (D). This observation suggests a mechanistic link between NSD2 activity and the turnover kinetics of H3.3, possibly implicating NSD2-mediated histone methylation in the regulation of H3.3 stability and turnover.

Additionally, our findings revealed that the loss of HIRA resulted in delayed decay not only of H3.3 but also of H3.1. This outcome suggests a broader role for HIRA beyond its established function in depositing H3.3 onto active chromatin. It implies that HIRA may also influence the turnover of both H3.1 and H3.3 histone variants, possibly through its involvement in chromatin remodeling or histone exchange processes (as illustrated in Fig. 5E), with detailed quantification provided in (F).

Overall, these results highlight the intricate interplay between histone-modifying enzymes such as NSD2 and chromatin remodeling factors like HIRA in orchestrating the dynamic turnover of histone variants. Further mechanistic studies are warranted to elucidate the precise molecular mechanisms underlying the observed effects of NSD2 and HIRA loss on H3.1 and H3.3 decay kinetics, which may have implications for understanding epigenetic regulation in various biological processes.

## Discussion

By conducting FRAP and live cell time-lapse imaging analysis, we gained valuable insights into the behavior of H3.3 within the nuclei of live mouse fibroblast cells. Our results revealed that the majority of H3.3 displayed high mobility, indicating its dynamic nature within the chromatin landscape. Additionally, we observed a smaller pool of H3.3 that exhibited less mobility.

Mathematical fitting of the FRAP data confirmed these observations and demonstrated that the behavior of H3.3 aligns best with a three-binding-state model, consisting of Mobile-Fast, Mobile-Slow, and Immobile (bound) fractions(37). Based on our estimates, approximately 50% and 28% of the preexisting populations of H3.3 transiently interacts with chromatin (i.e., Mobile-fast, and Mobile-slow) and while 23% remained more stably bound(immobile).

In contrast, H3.1 exhibited slower mobility compared to H3.3, with approximately 74% of the preexisting H3.1 fraction being stably bound (immobile), while 21% and 5% represented Mobile-Fast and Mobile-Fast fractions exchanging with chromatin, respectively. The recovery curve of H3.1 displayed dramatically reduced mobility (74%), suggesting most of H3.1 being chromatin bound most of time. On the other hand, the mobility of preexisting H3.3 appeared to correlate with its transcriptional competence, indicating its ability to interact with chromatin. The more stably bound species of H3.3 observed in FRAP may partially represent its binding to chromatinized targets within live nuclei(37,38). Interestingly, even the mobile-slow species of H3.3(28%) displayed faster recovery than H3.1(5%), suggesting that this species also interacts with targets to some degree, although some of them may be nonspecifically scanning chromatin. The FRAP measurements of H3.3 mobility primarily indicated a steady-state, genome-wide interaction with chromatin. Regarding newly synthesized H3.3, most of the newly synthesized fraction was found to be chromatin bound (immobile fraction), with the remaining fractions exhibiting fast (29%) and slow (11%) exchange with chromatin. These results suggest that newly synthesized H3.3 also follows a three-phase binding kinetics, similar to the preexisting fraction of H3.3.

Furthermore, we observed significant alterations in the recovery profile of H3.3 when cells were exposed to IFNβ. This included a delay in the initial phase of H3.3 recovery and an increase in the less mobile component (bound immobile fraction). A similar trend was observed in the newly synthesized fraction of H3.3. These findings indicate that IFNβ stimulation leads to an increased genome-wide interaction of H3.3 with chromatin during ongoing transcription. Thus, the signal-induced shift in mobility likely represents a global modification of the interaction between H3.3 and chromatin, which is likely linked to changes in gene expression. However, the precise mechanism by which IFNβ alters H3.3 mobility remains mostly unclear(18,39).

Moreover, our experiments involving transcriptional inhibition by Actinomycin D and Flavopiridol revealed that both compounds impeded the mobility of H3.3. This pattern was also observed in the newly synthesized fraction of H3.3, suggesting their direct involvement in transcriptional processes.

Loss of HIRA and NSD2 further supported the importance of these proteins for H3.3 mobility. The recovery profile of H3.3 in the absence of HIRA and NSD2 displayed an additive effect, indicating that these two partners can independently and jointly influence H3.3 mobility. This observation is consistent with previous reports suggesting the formation of a ternary complex involving H3.3, HIRA, and NSD2 in certain cases(18).

Histone turnover more specifically decay is a fundamental process in chromatin dynamics, influencing gene expression patterns and cellular functions. Our investigation into the decay dynamics of H3.1 and H3.3 histone variants provides valuable insights into the regulation of chromatin structure and function(28,40).

The distinct roles of H3.1 and H3.3 in chromatin regulation have been highlighted in our study. H3.1, associated with constitutively active chromatin regions, and H3.3, preferentially deposited at actively transcribed sites, exhibit differential decay kinetics. The observed differences in decay rates between H3.1 and H3.3 underscore their unique regulatory mechanisms and functional significance in chromatin dynamics.

Transcriptional inhibition shed light on the interplay between transcriptional activity and histone turnover. The delayed decay of H3.3 upon transcriptional inhibition suggests a dynamic response of histone turnover to changes in transcriptional programs. This observation underscores the dynamic nature of chromatin remodeling and its responsiveness to cellular cues.

Our study also elucidates the roles of histone-modifying enzymes, NSD2 and HIRA chaperone, in regulating histone turnover dynamics. Loss of NSD2 specifically impacts the decay of H3.3, implicating NSD2-mediated histone methylation in H3.3 stability. Conversely, loss of HIRA delays the decay of both H3.1 and H3.3, suggesting a broader role for HIRA in histone turnover regulation beyond its canonical function in H3.3 deposition.

Understanding the dynamics of histone turnover has significant implications for epigenetic regulation and disease pathogenesis. Dysregulation of histone turnover has been implicated in various pathological conditions, including cancer and neurodegenerative disorders.

In summary, our study offers invaluable insights into the dynamic behavior of H3.3 within live mouse fibroblast cells. Our findings illuminate the continuous interaction of H3.3 with chromatin across the nucleus, showcasing its distinct binding kinetics and reaffirming its direct involvement in transcriptional processes. Notably, a substantial portion of H3.3 displays high mobility, while a smaller fraction exhibits reduced mobility, indicating potential regulatory roles. Furthermore, our exploration of the effects of transcriptional arrest on H3.3 decay enriches our understanding of its spatiotemporal dynamics, providing deeper insights into its regulatory mechanisms.

These findings enhance our understanding of the intricate mechanisms underlying gene expression control and highlight potential avenues for future research.

## Supporting information

FRAP_supplementary_figures_bioRxiv.pdf 2.zip

## Authors contributions

V.N. performed all the experiments. V.N., D.B., T.S.K., and A.M. were involved in conducting the FRAP and timelapse videos analyses. V.N. did Bioinformatic analysis and drafted the manuscript. D.K., A.C., T.S.K and K.O. provided valuable input and guidance during the experiments and reviewed the manuscript. K.O. provided project oversight and direction. V.N. and K.O. finalized the draft.

## Data availability

The data supporting the findings of this study are available within the manuscript.

## Supporting information

This article contains supporting information.

## Acknowledgments

We thank Carl Wu for valuable discussion and constructive experimental feedback. We would like to acknowledge James Russell, Lynne Karycki Holtzclaw, and Vincent Schram of the Microscopy and Imaging Core (MIC)/NICHD/NIH for their initial assistance and guidance in operating the spinning disk microscope (Nikon) for acquiring time-lapse videos. Additionally, we extend our thanks to Peter Adams for generously sharing the anti-HIRA hybridoma cells/clones. We are also grateful to the members of the Division of Developmental Biology (DDB) and the Ozato lab for their valuable advice. This research made use of the computational resources provided by the National Institutes of Health High Performing Computation (HPC) Biowulf cluster.

