## Supplementary material for "Dynamic and decay kinetics of H3 variants in live cells reveal the pivotal role of HIRA/NSD2 in maintaining the distinct H3.3 specific chromatin landscape": FRAP_supplementary_figures_bioRxiv.pdf 2.zip: FRAP_supplementary_figures_bioRxiv.pdf

Nehru V et al.

### Supplementary Figures and Tables

#### Figure Highlights:

**Supplementary Figure. S1**. Cell lines expressing SNAP-tag with Histone H3 variants

**Supplementary Figure. S2**: Compared to global H3.3, newly synthesized H3.3 is less dynamic even after IFNβ stimulation

**Supplementary Figure. S3**: NSD2 and HIRA are important for the dynamics of newly synthesized H3.3

**Supplementary Figure. S4:** Inhibition of transcription machinery affects newly synthesized H3.3 dynamics

**Supplementary Figure. S5A.** Methodology to extract and calculate different fractions(i.e., Mobile-fast, Mobile-slow and Immobile) and half-time.

**Supplementary Figure. S5B(continued).** Methodology to extract and calculate different fractions(i.e., Mobile-fast, Mobile-slow and Immobile) and half-time.

**Supplementary Figure. S6A-S6F.** Time-lapse videos show decay over 50h time period under different experimental conditions.

**Supplementary Figure. S7A-S7F.** Time-lapse videos shows decay upto 20h time period in presence of Flavopiridol (100 nM) under different experimental conditions.

**Supplementary Figure S8.** Altered decay kinetics of H3.1 exacerbated by the transcription inhibition and loss of NSD2/HIRA.

**Supplementary Table. T1:** Summarized Global H3.1 and H3.3 fractions

**Supplementary Table.T2:** Summarized Newly synthesized H3.3 fractions

**Supplementary Figure S1.** Cell lines expressing SNAP-tag with Histone H3 variants

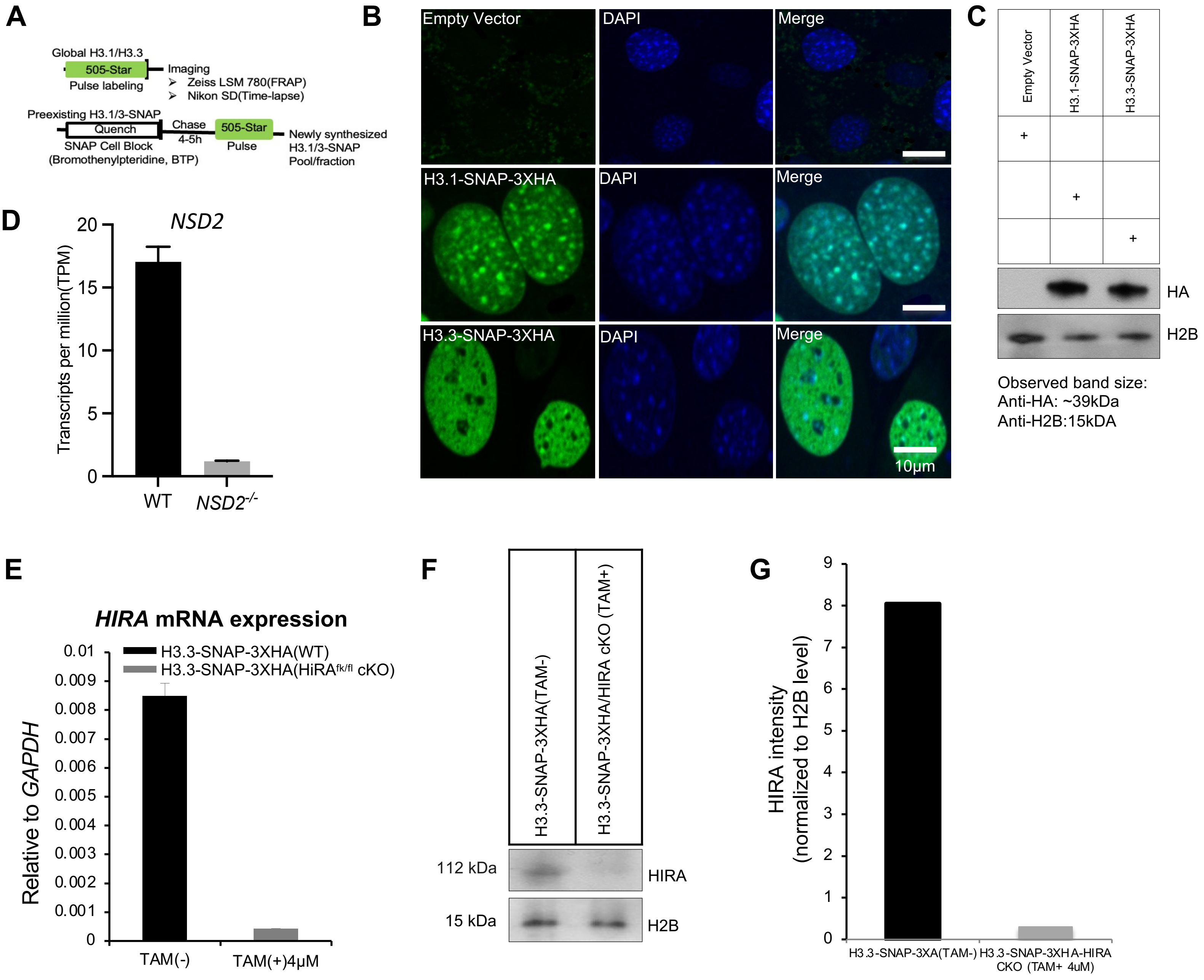

**Supplementary Figure S1. Cell lines expressing SNAP-tag with Histone H3 variants.**

**A.** The experimental scheme. Cell/nuclei permeable SNAP Cell 505-star dye (6-carboxyrhodamine 110, excitation maximum of 504 nm and an emission maximum of 532 nm) labelling of SNAP-tag stably co-expressing H3 variants, H3.1-SNAP-3XHA(WT) and H3.3-SNAP-3XHA(WT) in MEF cells.

**B.** SNAP (505-Star) labelling reveal H3.3-SNAP more euchromatic compared to H3.1-SNAP. Scale bars, 10 µm.

**C.** Expression of H3.1-SNAP-3XHA(~38KDa) and H3.3-SNAP-3XHA (~38KDa) detected in immunoblots from total protein extracts and probed with anti-HA-tag antibody and anti-H2B as a loading control.

**D.** *NSD2* total transcript count identified in RNA-seq analysis using Salmon suite/tool in H3.3-SNAP-3XHA(WT) and H3.3-SNAP-3XHA(*NSD2*$^{-/-}$) MEF cells at time 0h. (E) HIRA relative mRNA expression tested in both Wild-type (WT) and HIRA $^{fl/fl}$ cKO MEF cells after Tamoxifen(4uM) treatment for 4 days. The error bars represent standard error of the mean (SEM).

**F** and **G**. Expression and quantitation of Hira protein detected in immunoblots of the total proteins extracted from Wild-type (WT) and HIRA $^{fl/fl}$ Cre ERT MEF cells, probed with anti-Hira antibody (WC119) and anti-H2B as a loading control. The experiments were repeated three times.

**Supplementary Figure. S2:** Compared to global H3.3, newly synthesized H3.3 is less dynamic even after IFNβ stimulation

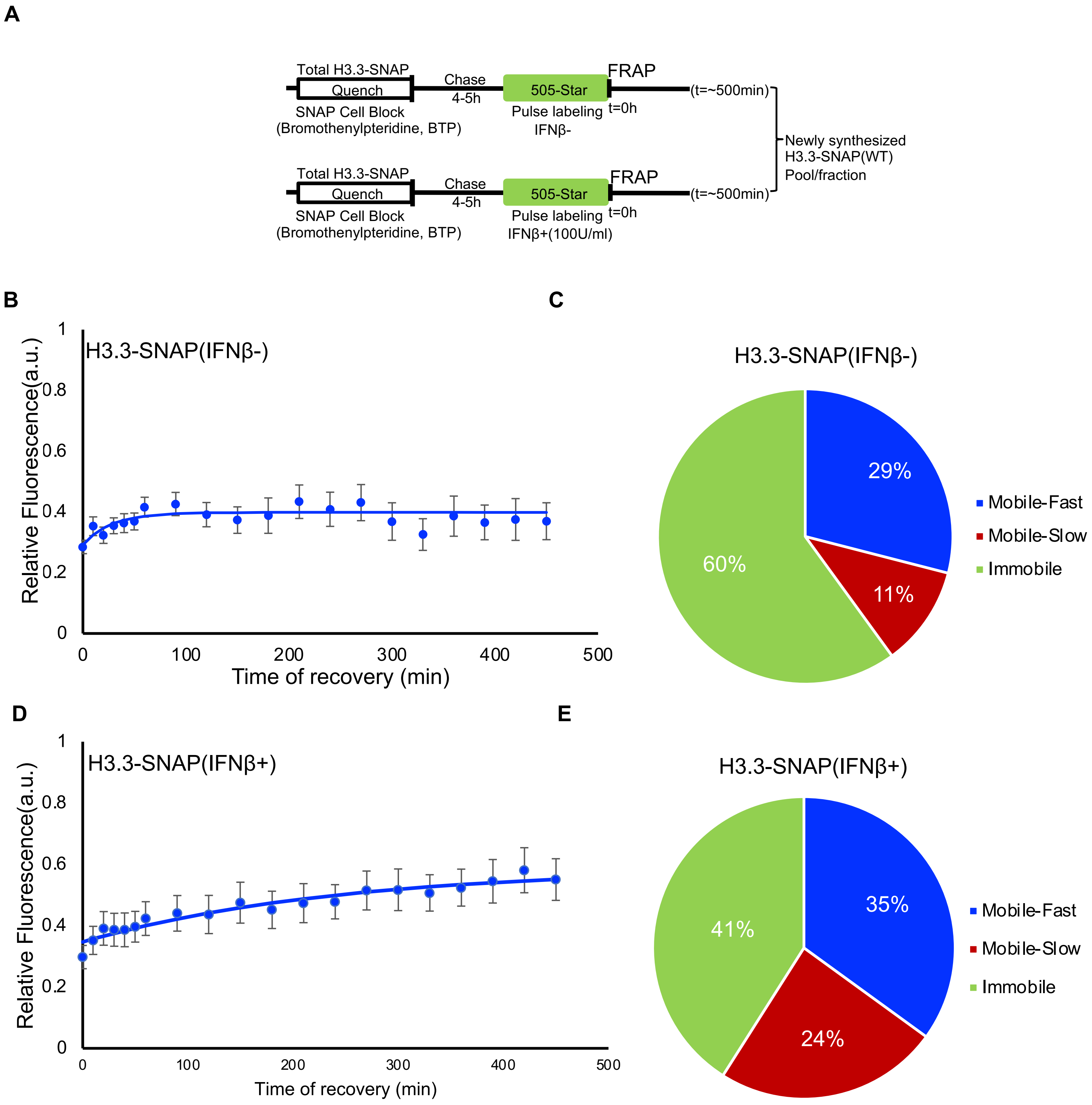

**Supplementary Figure. S2:** Compared to global H3.3, newly synthesized H3.3 is less dynamic even after IFNβ stimulation

**A.** Schematic representation of the *in vivo* SNAP quench-chase-pulse labeling strategy to detect newly synthesized H3.3 pool/fraction. At time $T_0$, the cells are incubated with a nonfluorescent SNAP substrate, BTP (quench) rendering the existing cellular pool of H3.3-SNAP pool unavailable for subsequent fluorescent labeling. After a period of substrate washout, referred to as the chase time (4 hours), the cells continue synthesizing H3.3-SNAP that remain unlabeled. Subsequently, at a specified chase time, the newly synthesized nascent proteins are selectively labeled with a fluorescent 505-Star (pulse) dye. This results in the generation of a distinct and fluorescent pool of SNAP-tagged proteins, representing the newly synthesized proteins or fraction. This nascent fluorescent pool was subjected to FRAP

**B** and **D**. FRAP curves show new H3.3 pool is less dynamic compared to global perhaps due to relatively less abundance. However, upon IFNβ stimulation, the new H3.3-SNAP(WT) pool becomes increasingly dynamic.

**C** and **E**. The labeled SNAP-tagged H3.3 protein is incorporated into chromatin through natural chromatin assembly mechanisms. The newly labeled H3.3 molecules replace the existing histone proteins within the nucleosomes, forming a pool of nucleosomes containing the newly synthesized H3.3. This pool of H3.3 was further analyzed to investigate its population percentage and associated half-life (t1/2) of different kinetic fractions that were determined using data from panels B and D and in Supplementary Table. T2B.

**Supplementary Figure.S3:** NSD2 and HIRA are important for the dynamics of newly synthesized H3.3

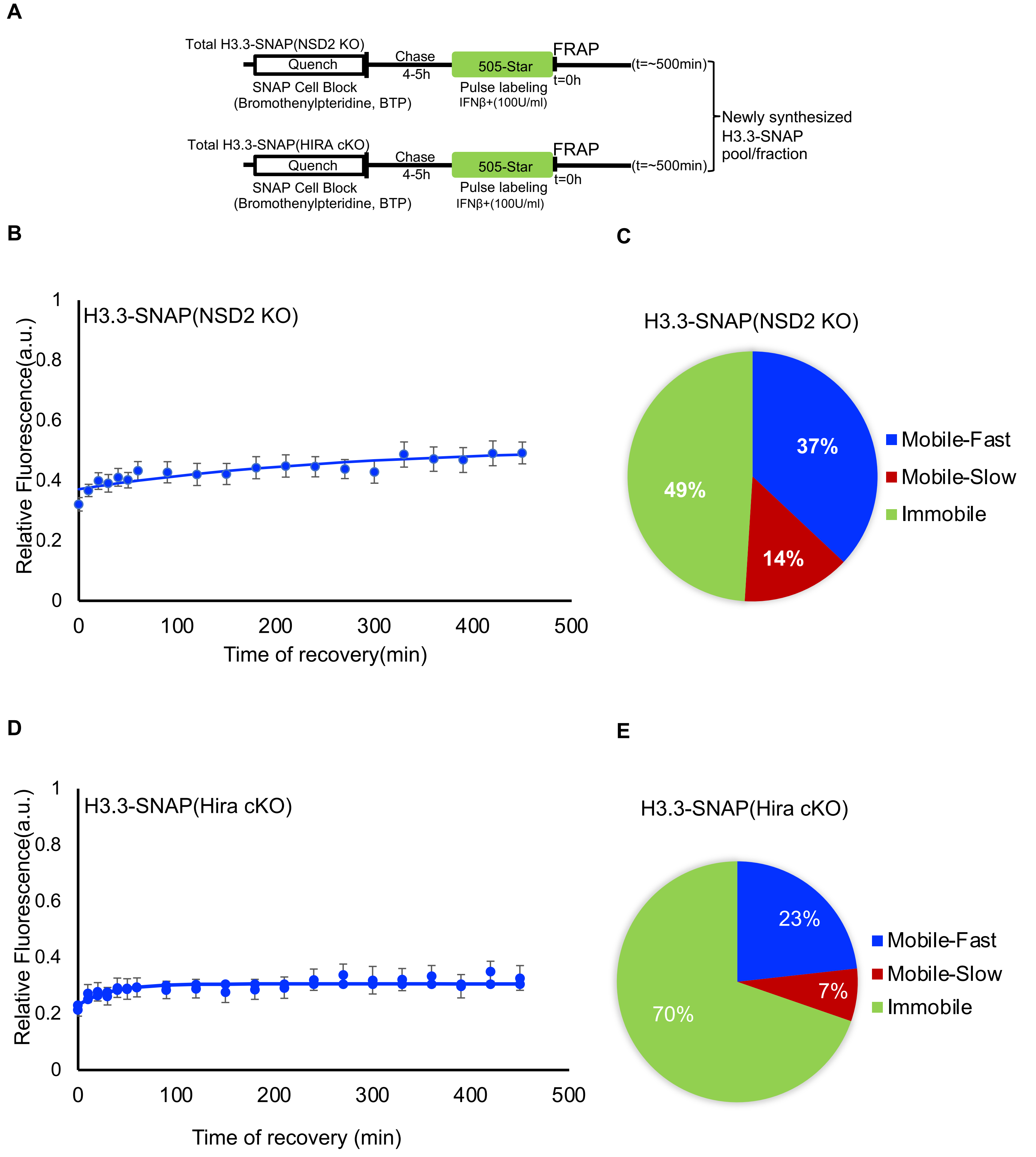




**Supplementary information**

**Supplementary Figure. S3** Impact of Hira and NSD2 on Dynamics of Newly Synthesized H3.3

**A**. Schematic representation illustrating the in vivo SNAP quench-chase-pulse labeling strategy employed to detect the dynamics of the newly synthesized H3.3 pool/fraction (described in Figure S3). The nascent fluorescent pool was subjected to FRAP analysis.

**B** and **D**. FRAP curves demonstrate the effect of HIRA and NSD2 loss on the mobility and exchange of the newly synthesized H3.3 pool.

**C** and **E**. Analysis reveals that HIRA loss leads to approximately 70% of the new H3.3 pool being chromatin bound or "immobile," while 23% and 7% exhibit "Mobile-Fast" and "Mobile-Slow" fractions, respectively. Conversely, loss of NSD2 results in 49% of the newly labeled H3.3 molecules being immobile, or chromatin bound, with 37% and 14% displaying "Mobile-Fast" and "Mobile-Slow" fractions, respectively.

These population percentages and associated half-life (t1/2) values for the different kinetic fractions were determined using data from panels B and D, as well as Supplementary Table. T2.

**Supplementary Figure. S4:** Inhibition of transcription machinery affects newly synthesized H3.3 dynamics

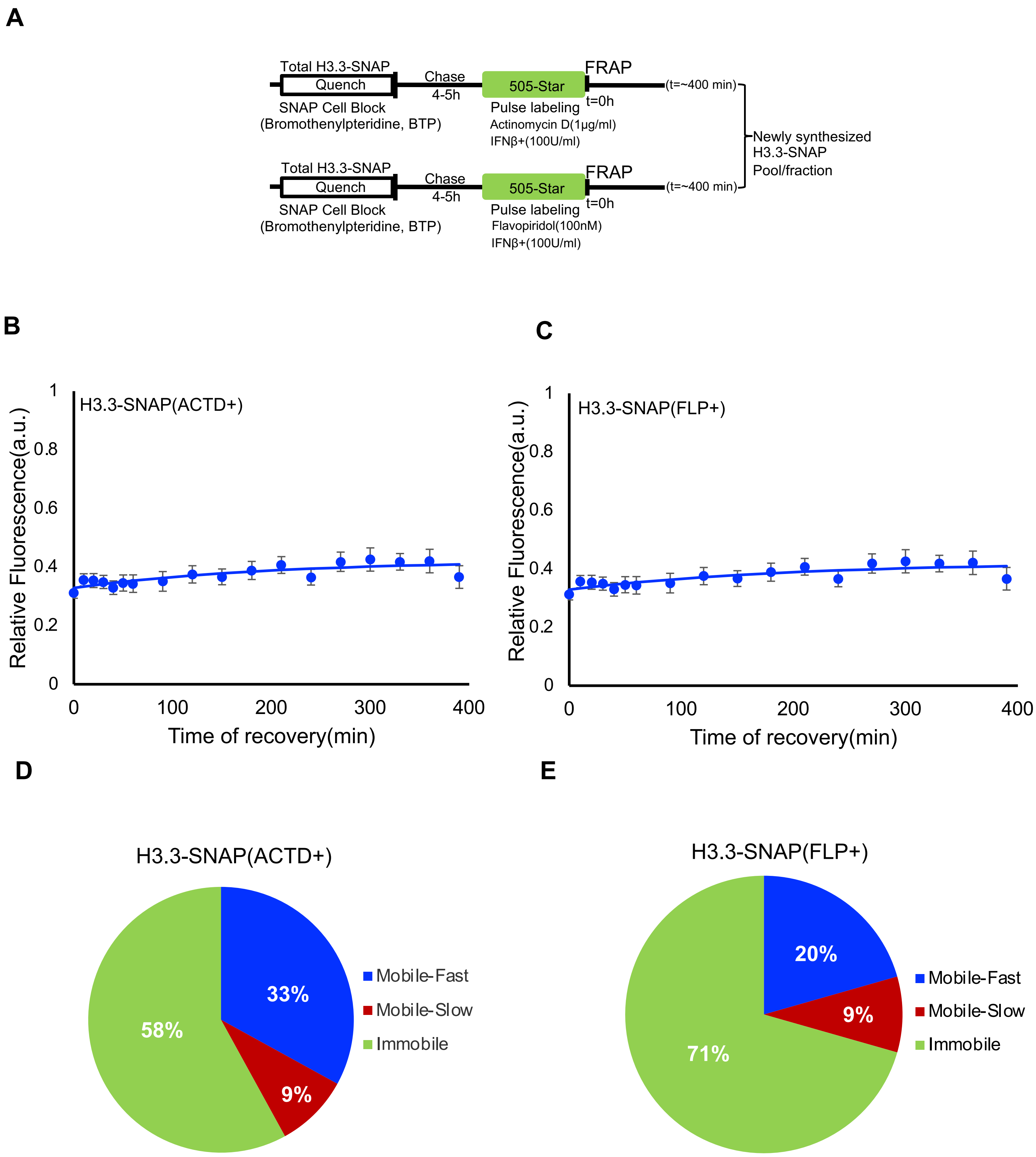

**Supplementary Figure. S5:** Inhibition of transcription machinery affects newly synthesized H3.3 dynamics

**A**. Schematic representation of the *in vivo* SNAP quench-chase-pulse labeling strategy to detect newly synthesized H3.3 pool/fraction. At time $T_0$, the cells are incubated with a nonfluorescent SNAP substrate block, Bromothenylpteridine (BTP) (quench) rendering the existing cellular pool of H3.3-SNAP pool unavailable for subsequent fluorescent labeling. After a period of substrate washout, referred to as the chase time (around 4 hours), the cells continue synthesizing H3.3-SNAP that remain unlabeled. Subsequently, at a specified chase time, the newly synthesized nascent proteins are selectively labeled with a fluorescent 505-Star (pulse) dye. This results in the generation of a distinct and fluorescent pool of SNAP-tagged proteins, representing the newly synthesized proteins or fraction. This nascent fluorescent pool was subjected to FRAP.

**B** and **C**. FRAP curves: Cell treated with Actinomycin D and Flavopiridol arrest (immobile chromatin bound fraction) newly synthesized H3.3 mobility even after IFNβ stimulation reinforcing H3.3 role in transcription couple deposition.

**D** and **E**. Population percentage and associated half-life (t1/2) of different kinetic fractions were determined using data from panels B-D and in Supplementary Table. T2. Actinomycin D(1ug/ml) and Flavopiridol(100mM) impede newly synthesized H3.3 mobility despite IFNβ treatment.

**Supplementary Figure. S5A**: Methodology to extract and calculate different fractions(i.e., Mobile-fast, Mobile-slow and Immobile) and half-time.

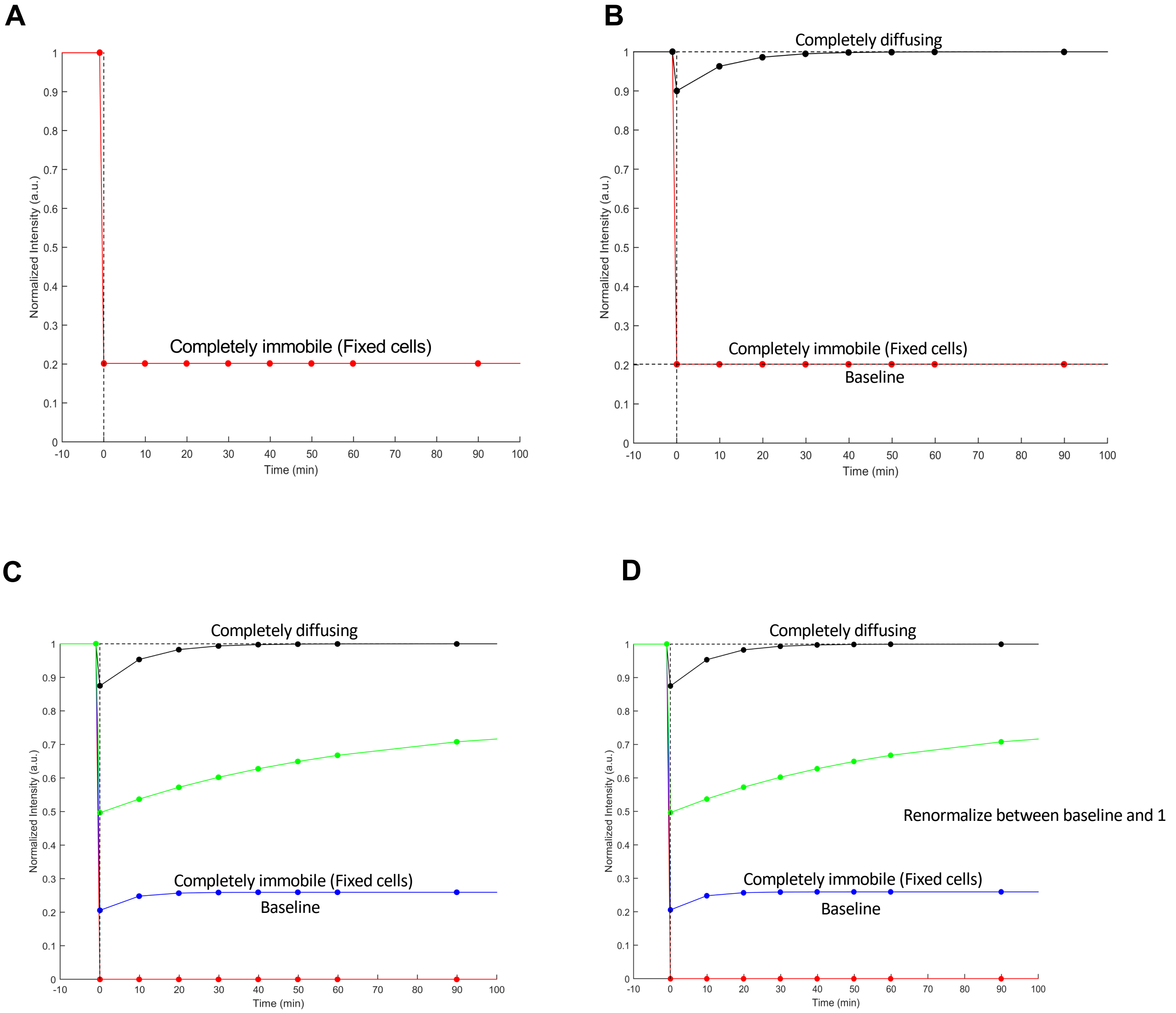

**Supplementary Figure. S5B(continued).** Methodology to extract and calculate different fractions(i.e., Mobile-fast, Mobile-slow and Immobile) and half-time.

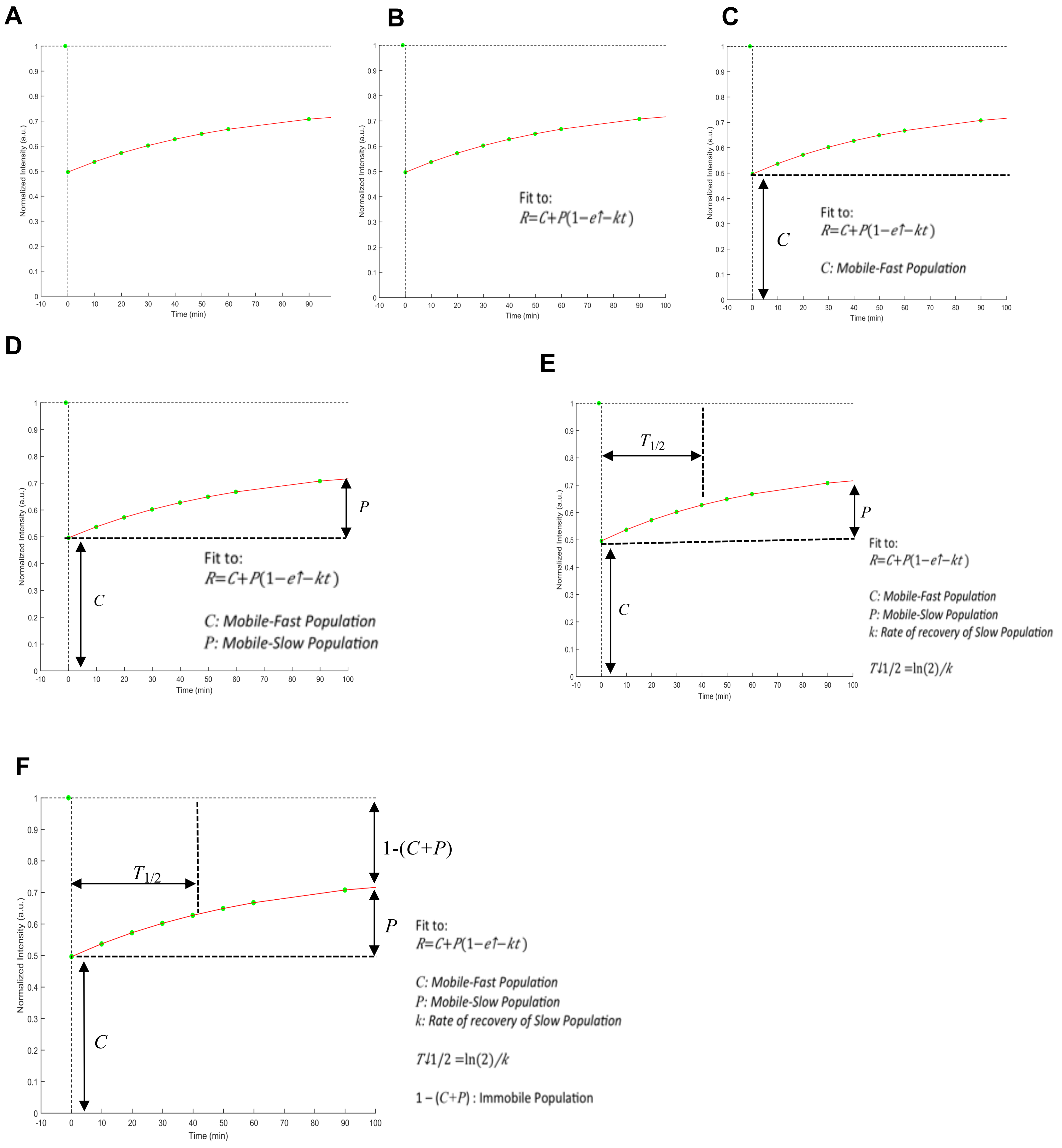

**Supplementary Figure. S5A and S5B.** Methodology to extract and calculate different fractions (i.e., Mobile-fast, Mobile-slow and Immobile).

S5A(A-D): Detailed methodology employed to extract and calculate the different kinetic fractions involved in chromatin dynamics analysis. The aim is to provide a comprehensive understanding of how the Mobile-fast, Mobile-slow, and Immobile fractions are determined.

S5B(A-F): Calculation of Mobile-fast, Mobile-slow, and Immobile fractions in chromatin dynamics analysis. This supplementary figure focuses on the calculation and quantification of the Mobile-fast, Mobile-slow, and Immobile fractions involved in the analysis of chromatin dynamics. It provides a detailed description of the equations, formulas, or algorithms used to determine the relative proportions of each fraction.

The figure may include illustrations or diagrams to visually explain the calculation process. It may also present representative data or examples to demonstrate the application of the methodology in practice.

Supplementary Figures. S5A and S5B collectively provide a comprehensive overview of the methodology employed to extract and calculate the different fractions and half-time values of H3.1-SNAP and H3.3-SNAP under different experimental conditions.

**Supplementary Figure. S6. Time-lapse shows decay over 50h time period**

**Supplementary Figure. S6A:** H3.1-SNAP-3XHA(WT)

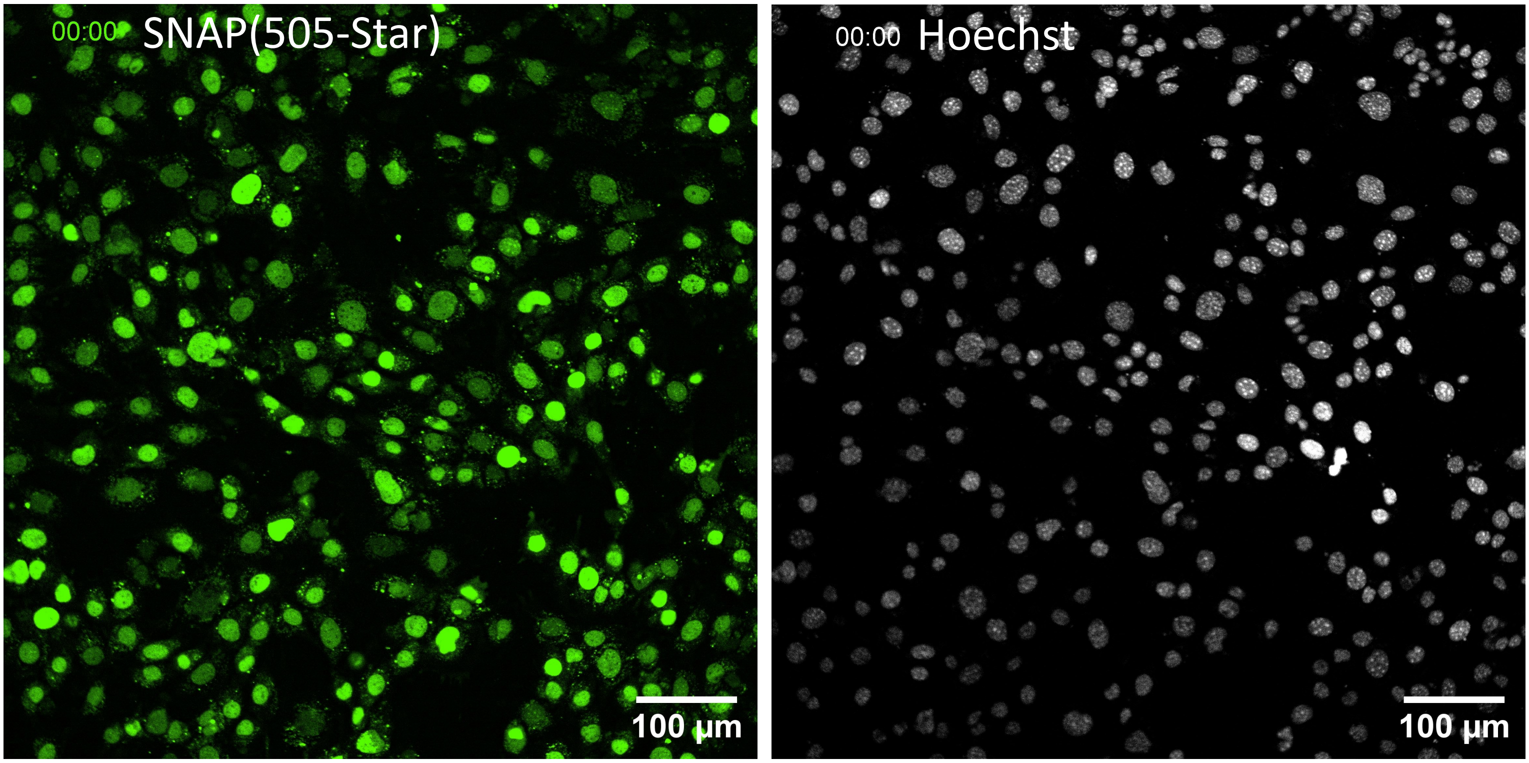


**Supplementary Figure. S6B:** H3.3-SNAP-3XHA(WT)

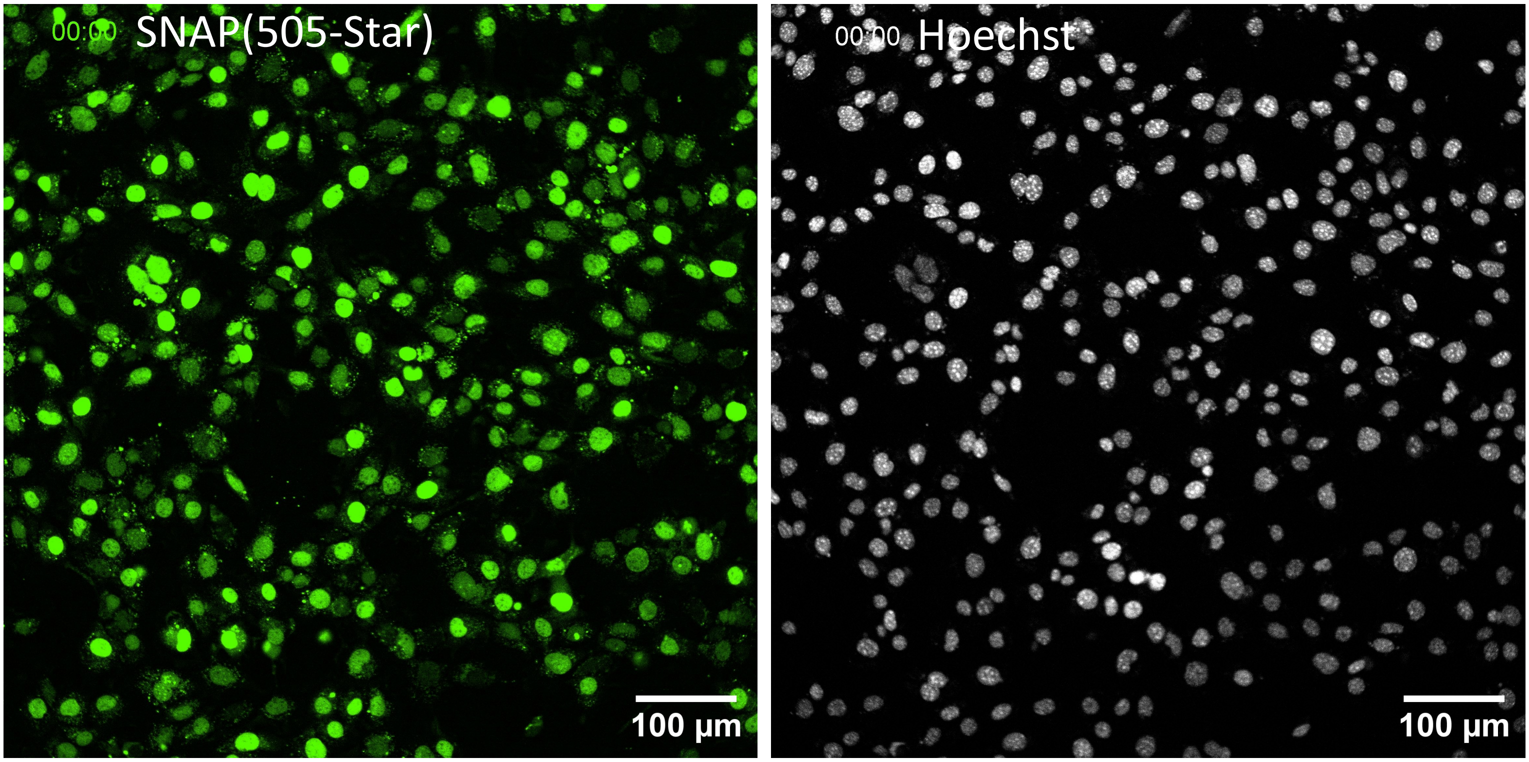

Supplementary Figure. S6(continued). Time-lapse shows decay over 50h time period

#### **Supplementary Figure. S6C:** H3.1-SNAP-3XHA(NSD2KO)

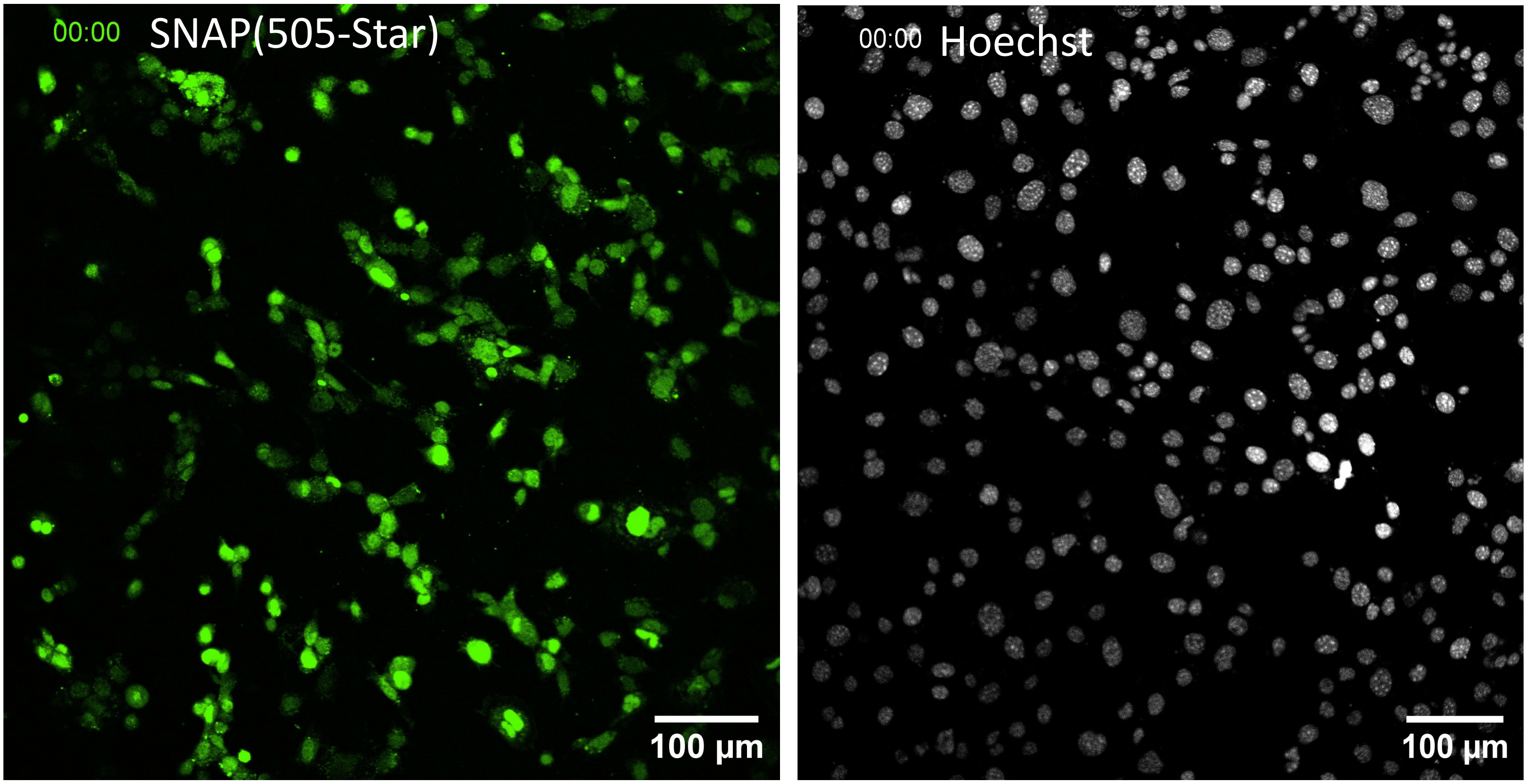


#### **Supplementary Figure. S6D:** H3.3-SNAP-3XHA(NSD2KO)

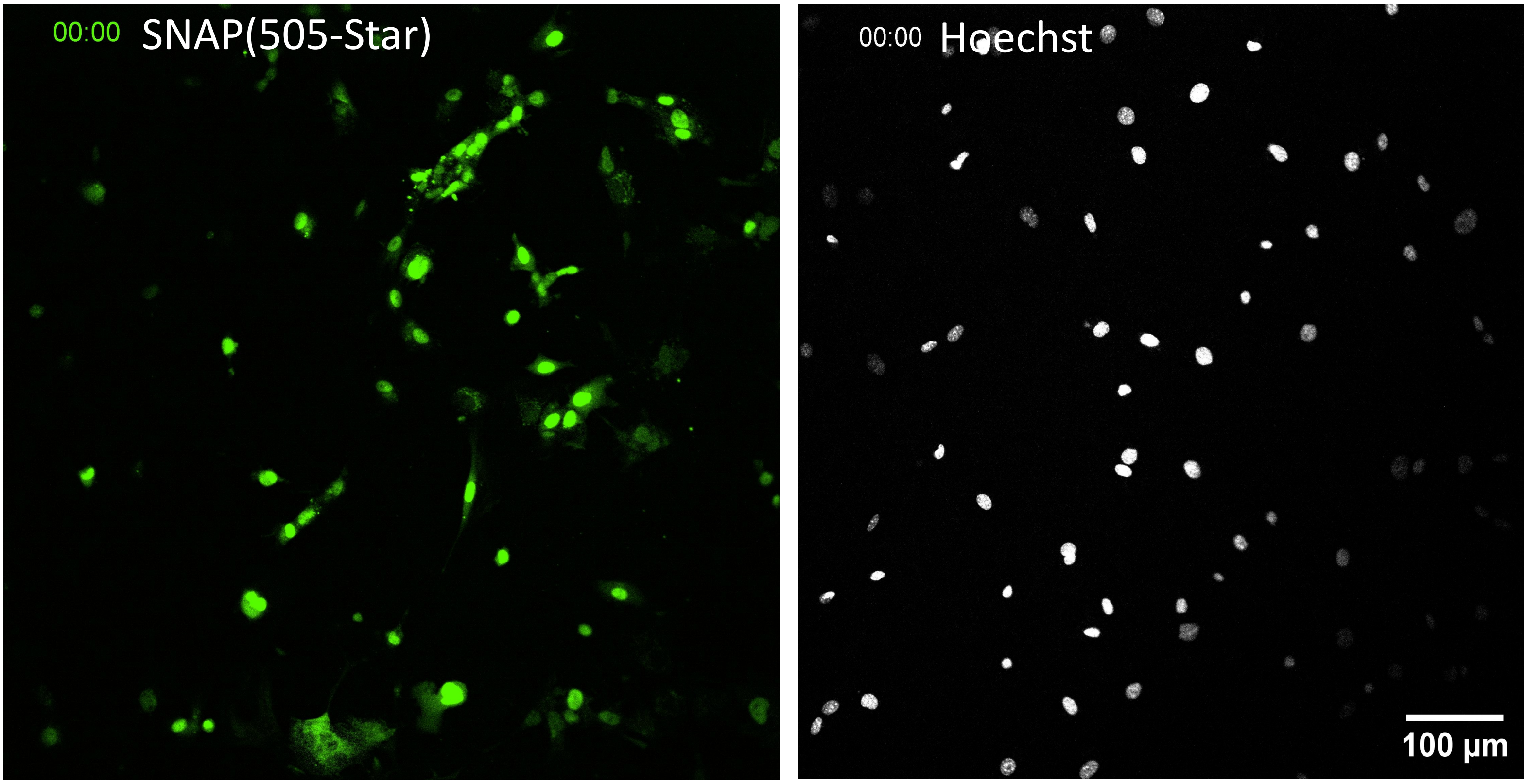

**Supplementary Figure. S6(continued). Time-lapse shows decay over 50h time period**

**Supplementary Figure. S6E:** H3.1-SNAP-3XHA(HIRA cKO)

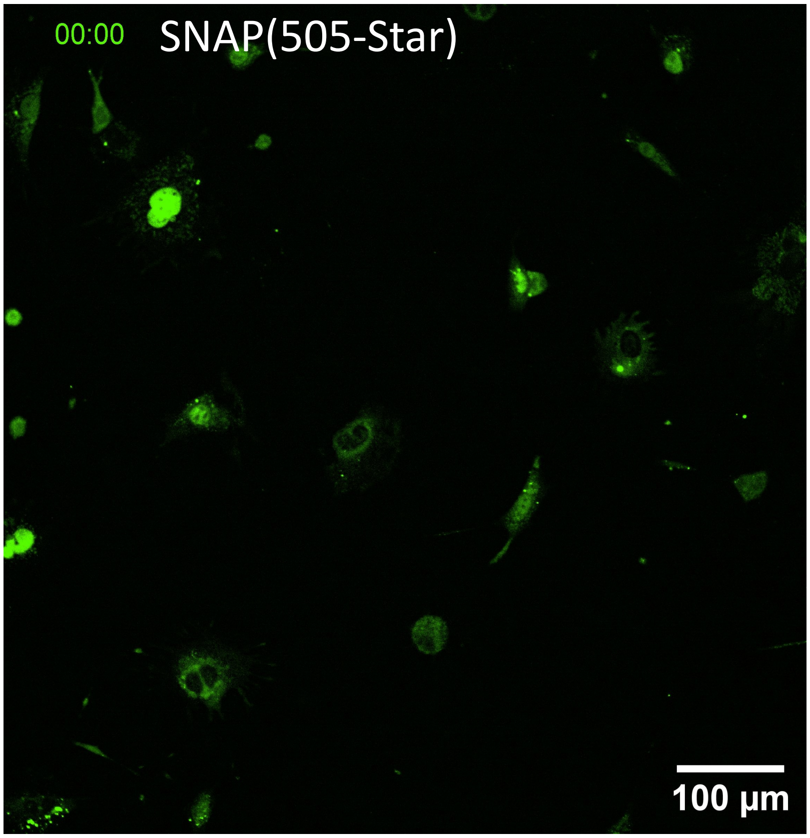


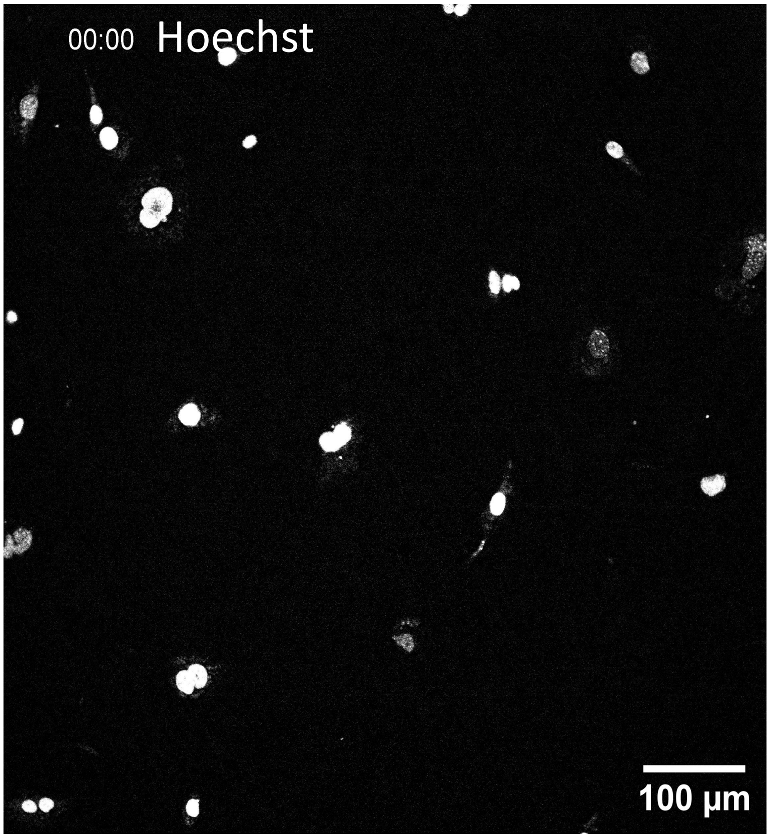


**Supplementary Figure. S6F:** H3.3-SNAP-3XHA(HIRA cKO)

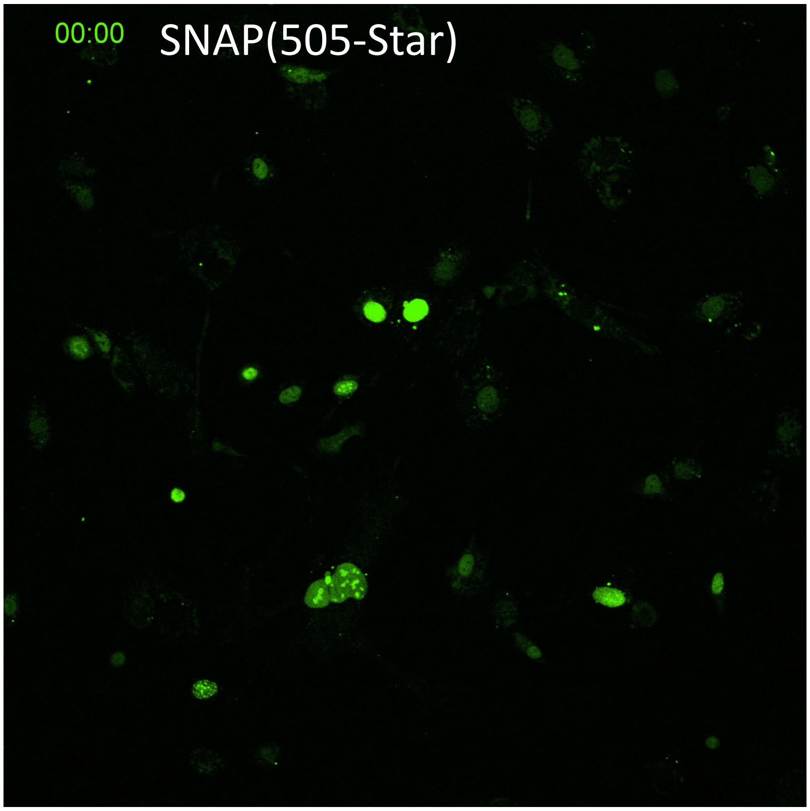


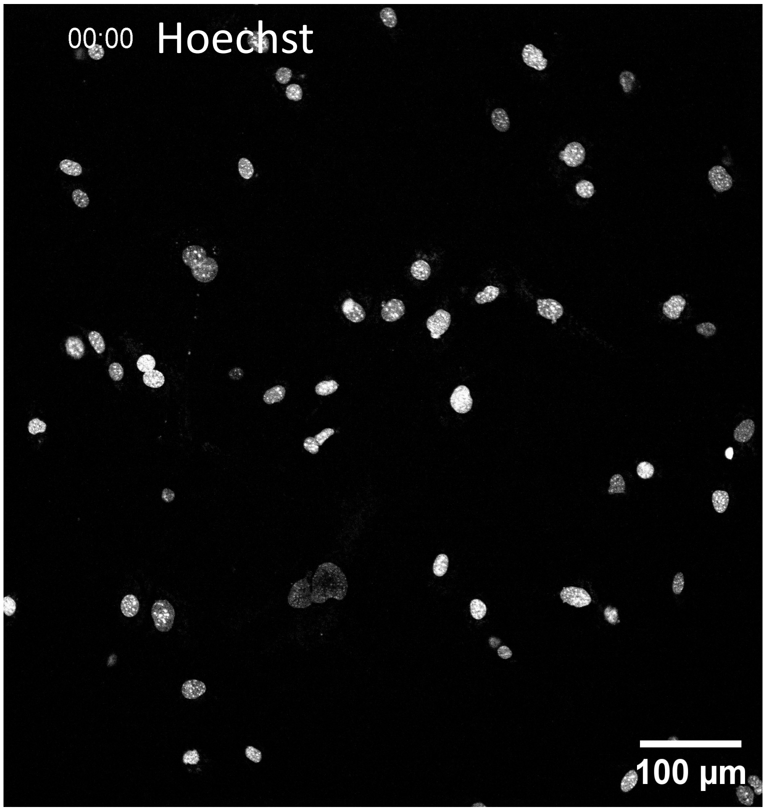




### Supplementary information

**Supplementary Figure. S6A.** Time-lapse video showing decay of H3.1-SNAP-3XHA(WT) over 50h time period

**Supplementary Figure. S6B.** Time-lapse video showing decay of H3.3-SNAP-3XHA(WT) over 50h time period

**Supplementary Figure. S6C.** Time-lapse video showing of H3.1-SNAP-3XHA(NSD2 KO) over 50h time period

**Supplementary Figure. S6D.** Time-lapse video showing of H3.3-SNAP-3XHA(NSD2 KO) over 50h time period

**Supplementary Figure. S6E.** Time-lapse video showing of H3.1-SNAP-3XHA(HIRA cKO) over 50h time period

**Supplementary Figure. S6F.** Time-lapse video showing of H3.3-SNAP-3XHA(HIRA cKO) over 50h time period

**Supplementary Figure. S7. Time-lapse shows decay over 20h time period in presence of Flavopiridol.**

#### **Supplementary Figure. S7A:** H3.1-SNAP-3XHA(WT)

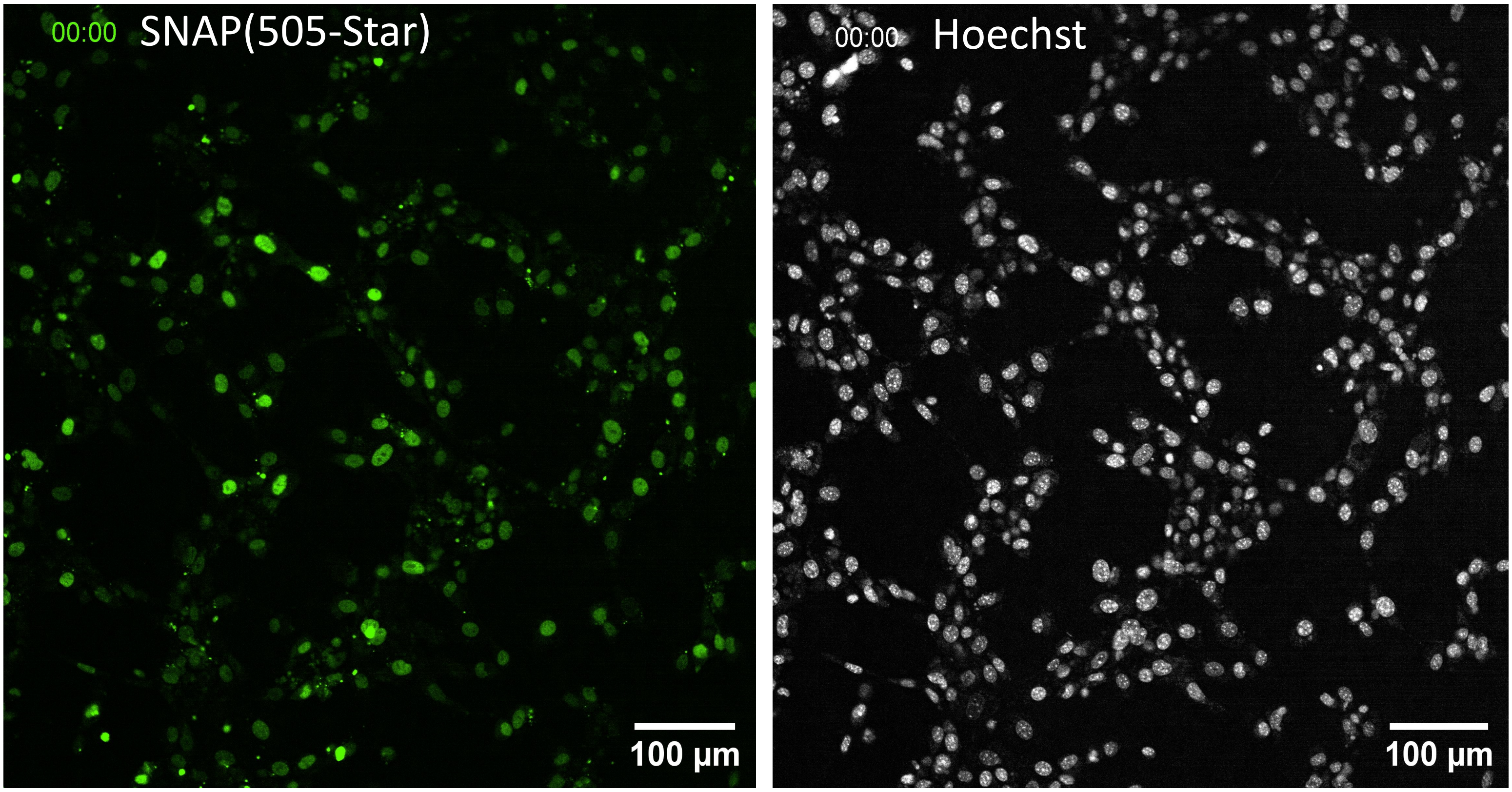


#### **Supplementary Figure. S7B:** H3.3-SNAP-3XHA(WT)

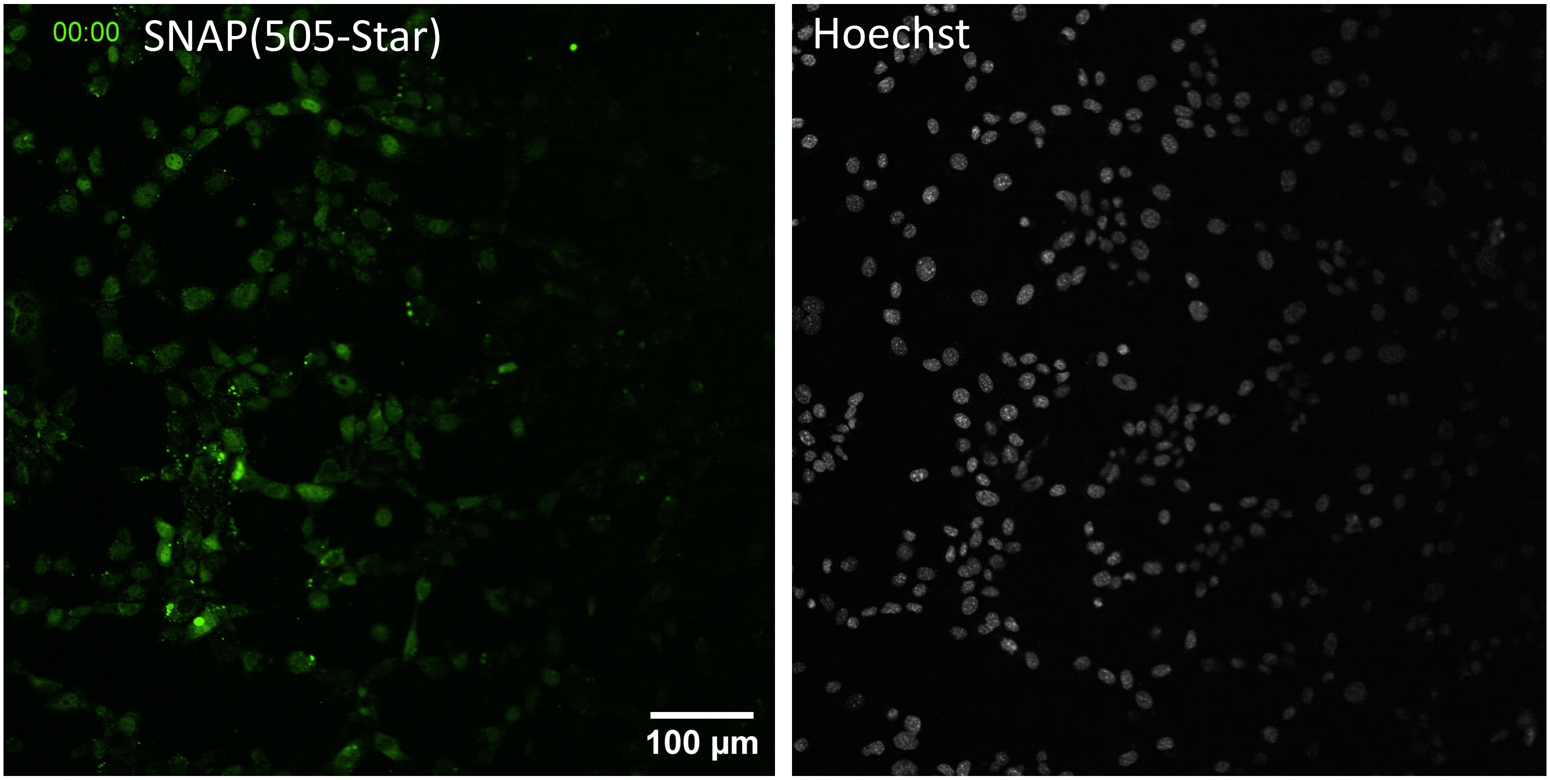

**Supplementary Figure. S7(continued). Time-lapse shows decay over 20h time period in presence of Flavopiridol(100nM)**

**Supplementary Figure. S7C:** H3.1-SNAP-3XHA(NSD2KO)

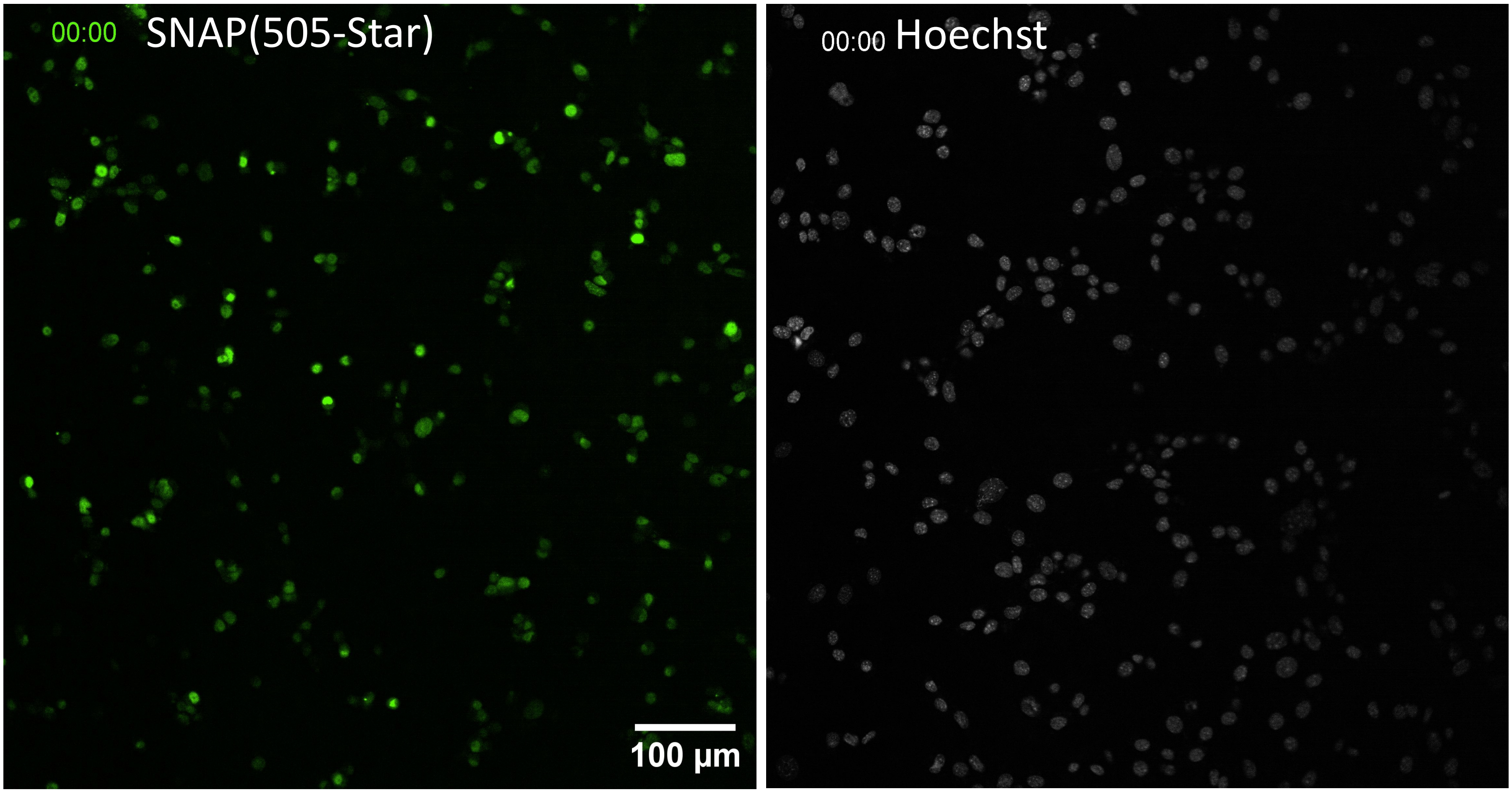


**Supplementary Figure. S7D:** H3.3-SNAP-3XHA(NSD2KO)

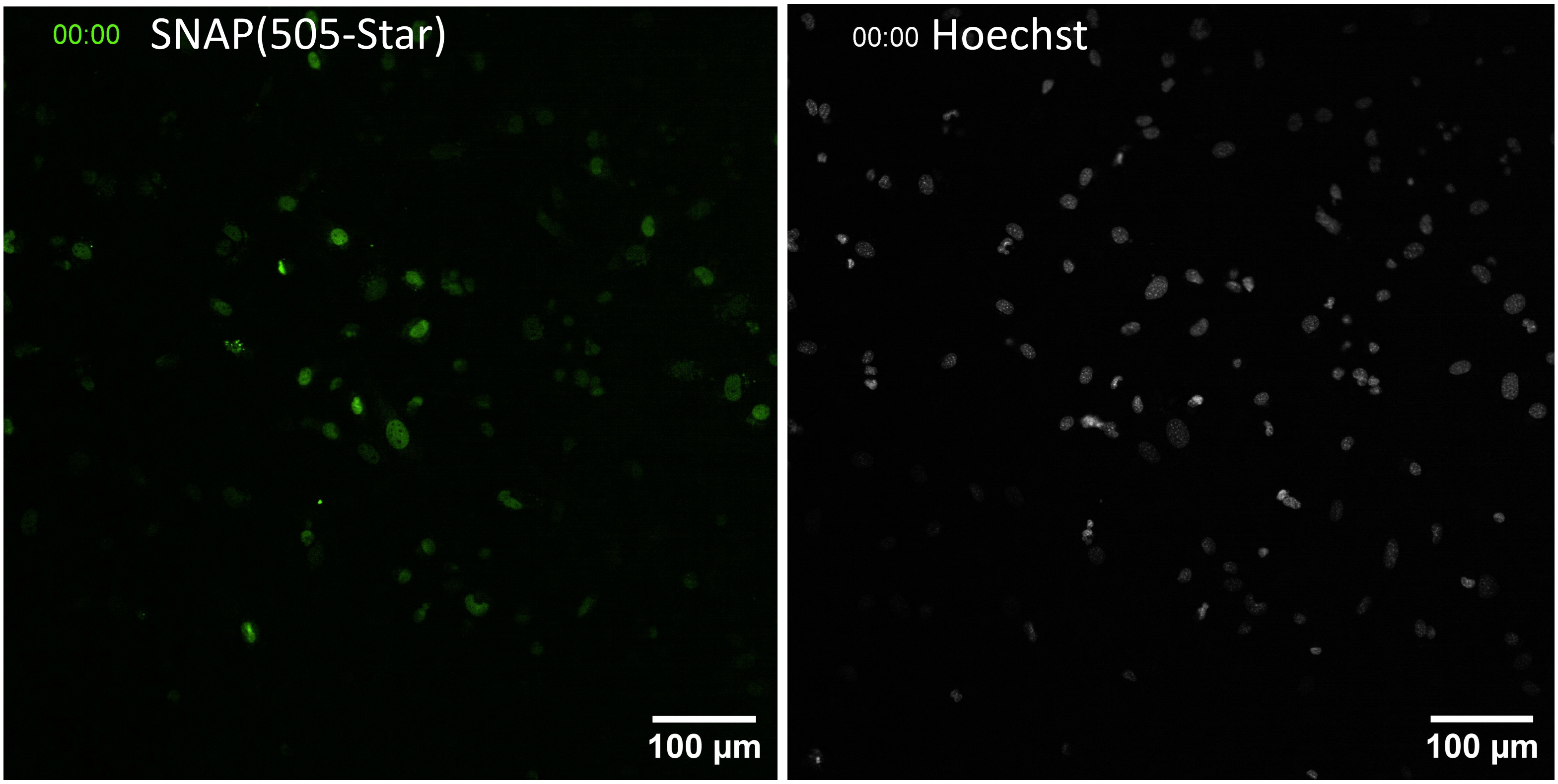




### Supplementary information

**Supplementary Figure. S7(continued). Time-lapse shows decay over 20h time period in presence of Flavopiridol.**

#### **Supplementary Figure. S7E:** H3.1-SNAP-3XHA(HIRA cKO)

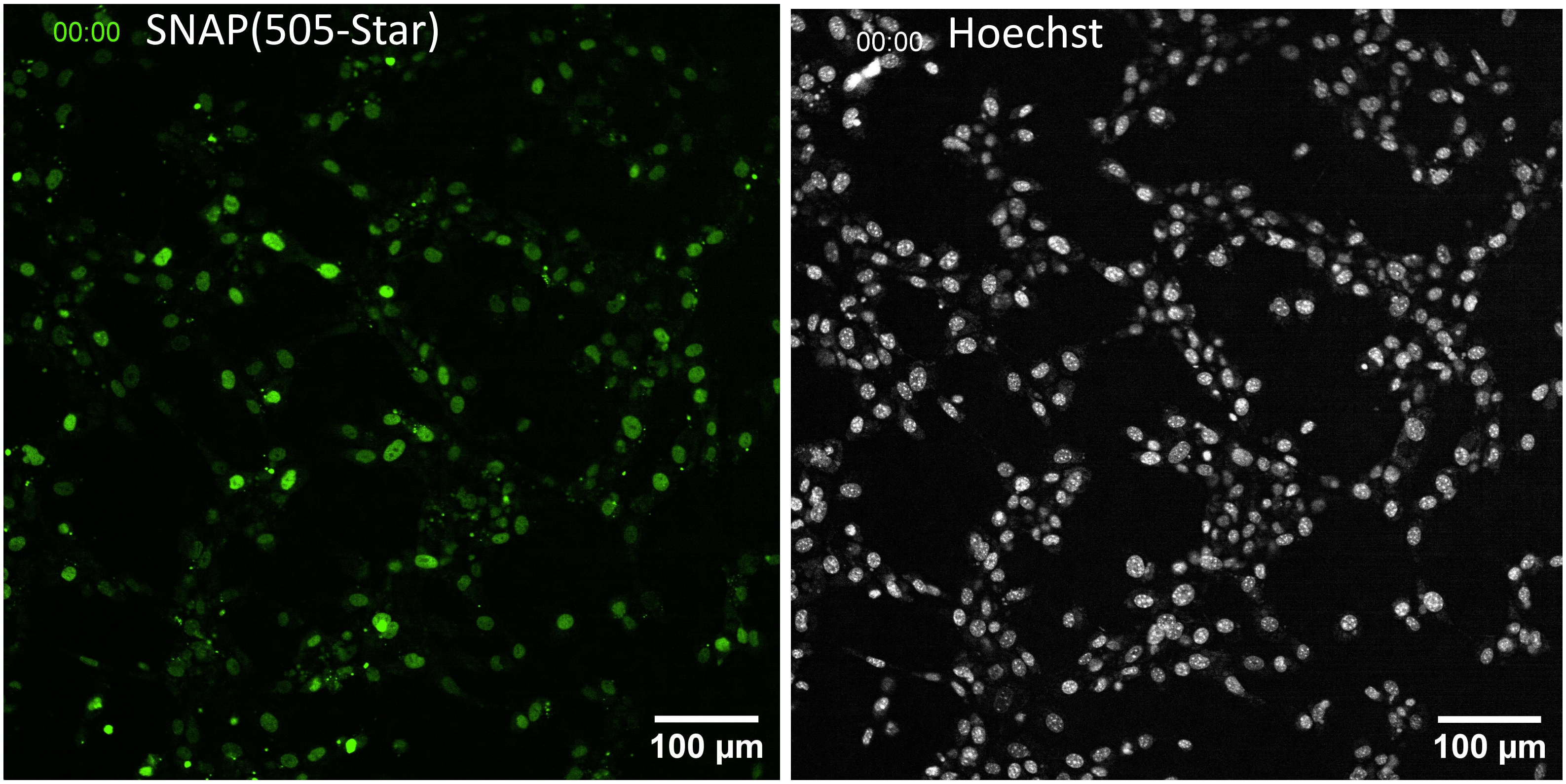


#### **Supplementary Figure. S7F:** H3.3-SNAP-3XHA(HIRA cKO)

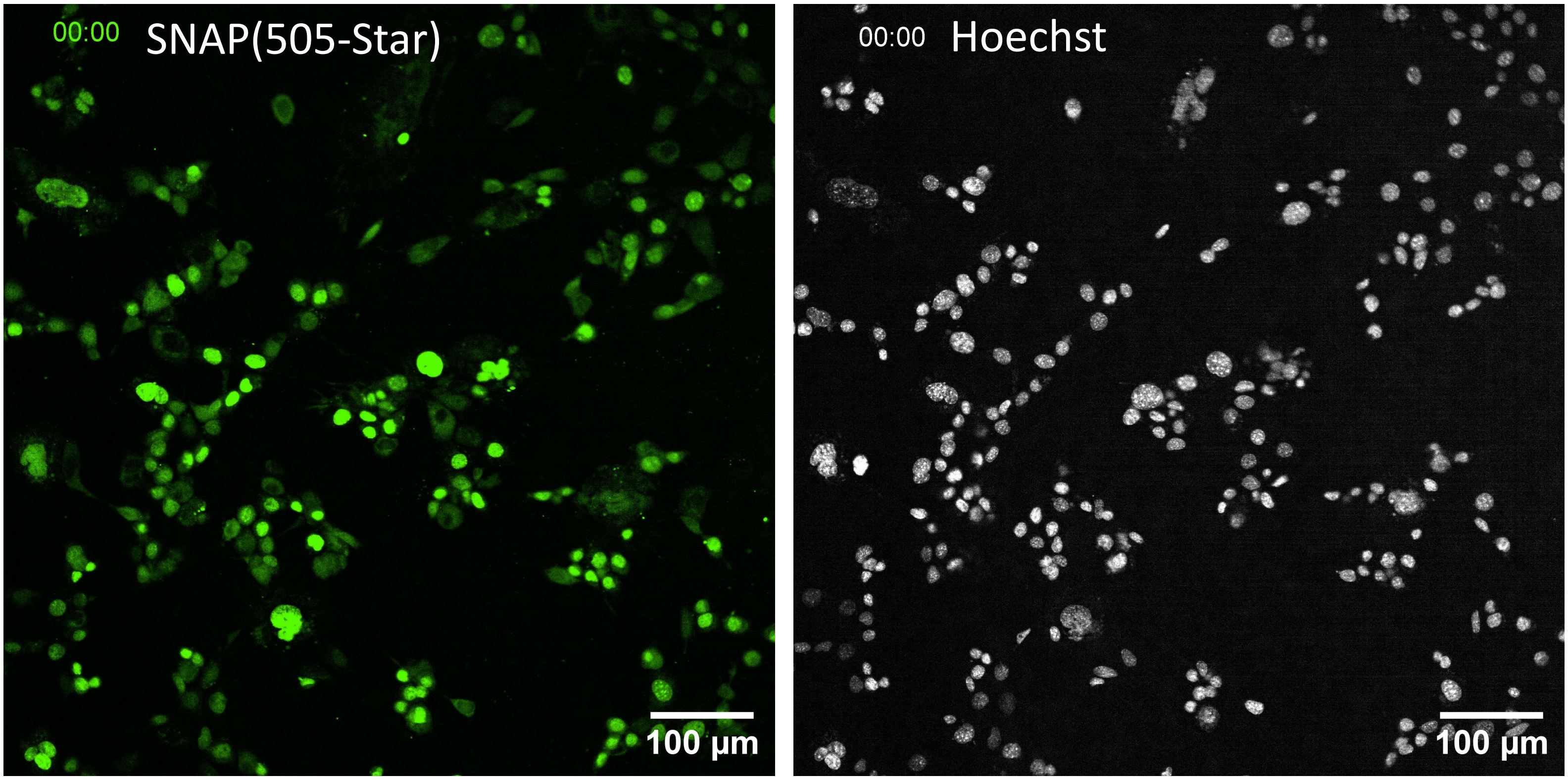

**Supplementary Figure. S7A.** Time-lapse video showing the decay of H3.1-SNAP-3XHA(WT) upto 20-hour period in the presence of Flavopiridol (100 nM).

**Supplementary Figure. S7B.** Time-lapse video showing the decay of H3.3-SNAP-3XHA(WT) upto 20-hour period in the presence of Flavopiridol (100 nM).

**Supplementary Figure. S7C.** Time-lapse video showing the decay of H3.1-SNAP-3XHA(NSD2 KO) upto 20-hour period in the presence of Flavopiridol (100 nM).

**Supplementary Figure. S7D.** Time-lapse video showing the decay of H3.3-SNAP-3XHA(NSD2 KO) upto 20-hour period in the presence of Flavopiridol (100 nM).

**Supplementary Figure. S7E.** Time-lapse video showing the decay of H3.1-SNAP-3XHA(HIRA cKO) upto 20-hour period in the presence of Flavopiridol (100 nM).

**Supplementary Figure. S7F.** Time-lapse video showing the decay of H3.3-SNAP-3XHA(HIRA cKO) upto 20-hour period in the presence of Flavopiridol (100 nM).

**Figure 8. Altered decay kinetics of H3.1 exacerbated by the transcription inhibition and loss of NSD2/HIRA.**

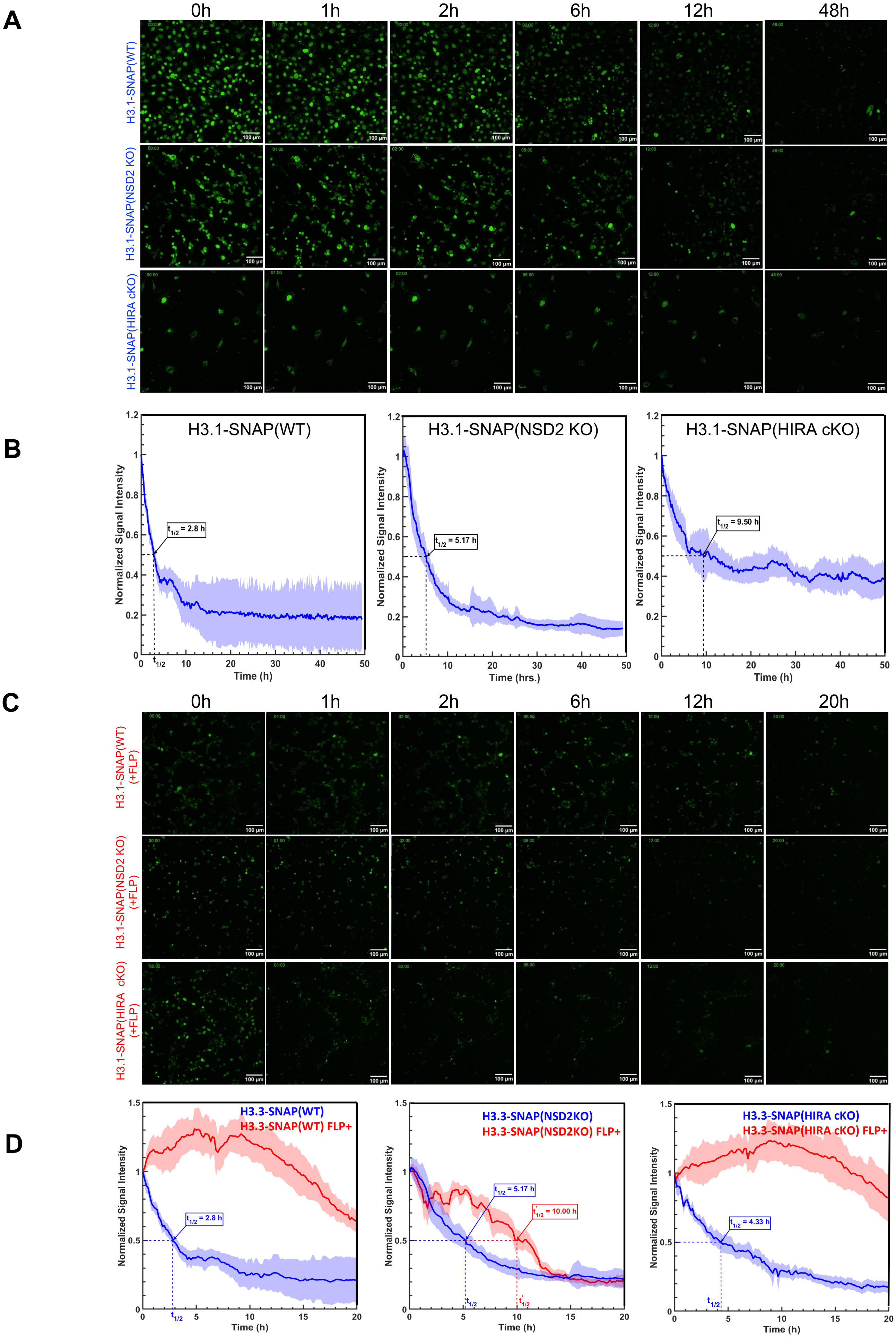




### Supplementary information

**Figure 8. Altered decay kinetics of H3.1 exacerbated by the transcription inhibition and loss of NSD2/HIRA.**

A. Displays the pulse-chase fluorescence intensity of H3.1-SNAP-3XHA (WT), H3.1-SNAP-3XHA (NSD2KO) and H3.1-SNAP-3XHA (HIRA cKO) at different timepoints (0 h, 1 h, 2h, 6h, 12h and 48 h).

B. Shows the fluorescence decay quantitation of H3.1-SNAP-3XHA (WT), H3.1-SNAP-3XHA (NSD2KO) and H3.1-SNAP-3XHA (HIRA cKO) (represented by the blue curves ) with $t^{1/2}$ values.

C. Displays the pulse-chase live-cell fluorescence intensity of H3.1-SNAP-3XHA (WT), H3.1-SNAP-3XHA (NSD2 KO), H3.1-SNAP-3XHA (HIRA cKO) at different timepoints (0 h, 1 h, 2h, 6h, 12h and 20 h) with Flavopiridol(100nM).

D. Shows the fluorescence decay quantitation of H3.1-SNAP-3XHA (WT) H3.1-SNAP-3XHA (NSD2KO) and H3.1-SNAP-3XHA (HIRA cKO) (represented by the red curve) with $t^{1/2}$ values with flavopiridol(100nM). All scale bars represent 100 μm.



#### Supplementary information

### Supplementary Table. T1A: Global H3.3 Fractions and Half-Time ($t^{1/2}$)

| Conditions (Global pool) | Mobile-fast fraction | Mobile-slow fraction | Half-time (min) | Immobile fraction | N |
|---|---|---|---|---|---|
| H3.1-SNAP-3XHA | 0.21 ± 0.03 | 0.05 ± 0.01 | 4.4 ± 10.4 | 0.74 ± 0.01 | 24 |
| H3.3-SNAP-3XHA | 0.50 ± 0.08 | 0.28 ± 0.08 | 43 ± 29 | 0.23 ± 0.12 | 18 |
| H3.3-SNAP-3XHA (Hira cKO) | 0.07 ± 0.03 | 0.07 ± 0.03 | 49 ± 60 | 0.86 ± 0.05 | 18 |
| H3.3-SNAP-3XHA (NSD2 KO) | 0.22 ± 0.03 | 0.28 ± 0.12 | 269 ± 240 | 0.50 ± 0.12 | 13 |
| H3.3-SNAP-3XHA(WT)-IFNβ+ (100U/ml) | 0.22 ± 2.6 | 0.78 ± 9.41 | 1433 ± 24000 | $1.3X10^{-5}$ ± 11.725 | 19 |
| H3.3-SNAP-3XHA (Hira cKO)-IFNβ+ (100U/ml) | 0.05 ± 0.03 | 0.06 ± 0.04 | 3.4 ± 6.9 | 0.89 ± 0.05 | 19 |
| H3.3-SNAP-3XHA(WT) Actinomycin D (1μg/ml) | 0.12 ± 0.04 | 0.20 ±0.04 | 75 ± 47 | 0.68 ± 0.07 | 18 |
| H3.3-SNAP-3XHA(WT) Flavopiridol (100nM) | 0.21 ± 0.03 | 0.18 ± 0.03 | 33 ± 12 | 0.61 ± 0.04 | 17 |
| H3.3-SNAP-3XHA(HIRA cKO) Actinomycin D (1μg/ml) | 0.10 ± 0.04 | 0.10 ±0.04 | 75 ± 47 | 0.90 ± 0.07 | 20 |
| H3.3-SNAP-3XHA(NSD2KO) Actinomycin D (1μg/ml) | 0.10 ± 0.04 | 0.10 ±0.04 | 75 ± 47 | 0.80 ± 0.07 | 20 |
| H3.3-SNAP-3XHA(HIRA cKO) Flavopiridol (100nM) | 0.15 ± 0.03 | 0.15 ± 0.03 | 33 ± 12 | 0.70 ± 0.04 | 17 |
| H3.3-SNAP-3XHA(NSD2 KO) Flavopiridol (100nM) | 0.20 ± 0.03 | 0.18 ± 0.03 | 20 ± 12 | 0.60 ± 0.04 | 17 |

### Supplementary Table.T2B: Newly Synthesized H3.3 Fractions and Half-Time ($t^{1/2}$)

| Conditions (New pool) | Mobile-fast fraction | Mobile-slow fraction | Half-time (min) | Immobile fraction | N |
|---|---|---|---|---|---|
| H3.3-SNAP-3XHA | 0.29 ± 0.07 | 0.11 ± 0.07 | 19.3 ± 24.1 | 0.60 ± 0.10 | 18 |
| H3.3-SNAP-3XHA(WT)-IFNβ+ (100U/ml) | 0.35 ± 0.02 | 0.24 ± 0.07 | 168 ± 119 | 0.41 ± 0.7 | 16 |
| H3.3-SNAP-3XHA(WT) Flavopiridol (100nM) | 0.20 ± 0.03 | 0.09 ± 0.03 | 26.2 ± 18.71 | 0.71 ± 0.04 | 19 |
| H3.3-SNAP-3XHA(WT) Actinomycin D (1ug/ml) | 0.32 ± 0.02 | 0.09 ± 0.07 | 137.7 ± 247.35 | 0.58 ± 0.07 | 16 |
| H3.3-SNAP-3XHA (Hira cKO) | 0.23 ± 0.05 | 0.07 ±0.05 | 22.0 ± 27.5 | 0.70 ± 0.07 | 11 |
| H3.3-SNAP-3XHA (Hira cKO)-IFNβ+ (100U/ml) | 0.20 ± 0.03 | 0.04± 0.03 | 26 ± 40 | 0.76 ± 0.4 | 15 |
| H3.3-SNAP-3XHA (NSD2 KO) | 0.37 ± 0.03 | 0.14 ± 0.06 | 187 ± 209 | 0.49 ± 0.07 | 13 |
| H3.3-SNAP-3XHA(HIRA cKO) Actinomycin D (1μg/ml) | 0.5 ± 0.04 | 0.5 ±0.04 | 75 ± 47 | 0.80 ± 0.07 | 10 |
| H3.3-SNAP-3XHA(NSD2 KO) Actinomycin D (1μg/ml) | 0.15 ± 0.04 | 0.15 ±0.04 | 75 ± 47 | 0.70 ± 0.07 | 10 |
| H3.3-SNAP-3XHA(HIRA cKO) Flavopiridol (100nM) | 0.10 ± 0.03 | 0.10 ± 0.03 | 33 ± 12 | 0.80 ± 0.04 | 17 |
| H3.3-SNAP-3XHA(NSD2 KO) Flavopiridol (100nM) | 0.20 ± 0.03 | 0.20 ± 0.03 | 20 ± 12 | 0.60 ± 0.04 | 17 |

**Supplementary Table. T1A: Global H3.1 and H3.3 Fractions**

Provides a comprehensive overview of the global H3.1 and H3.3 fractions and their corresponding half-time ($t^{1/2}$) values under different experimental conditions. It quantifies various kinetic fractions, including Mobile-fast, Mobile-slow, and Immobile, and represents the time required for half of each fraction to undergo turnover or exchange.

**Supplementary Table. T2B: Newly Synthesized H3.3 Fractions**

Shows fractions and half-time (t1/2) values of newly synthesized H3.3 under different experimental conditions. It provides a detailed analysis of the kinetic fractions involved in the turnover or exchange of the nascent H3.3 pool. The table includes the proportions of Mobile-fast, Mobile-slow, and Immobile fractions within the newly synthesized H3.3 pool, along with their corresponding half-time values.

Supplementary Tables T1A and T2B offer crucial quantitative information related to the dynamic properties of H3.3, providing insights into the turnover rates and relative contributions of different kinetic fractions in the dynamics of the global and newly synthesized H3.3 pool.
